# Aging impairs the essential contributions of non-glial progenitors to neurorepair in the dorsal telencephalon of the Killifish *N. furzeri*

**DOI:** 10.1101/2021.02.26.433041

**Authors:** Jolien Van houcke, Valerie Mariën, Caroline Zandecki, Sophie Vanhunsel, Lieve Moons, Rajagopal Ayana, Eve Seuntjens, Lutgarde Arckens

## Abstract

The aging central nervous system (CNS) of mammals displays progressive limited regenerative abilities. Recovery after loss of neurons is extremely restricted in the aged brain. Many research models fall short in recapitulating mammalian aging hallmarks or have an impractically long lifespan. We established a traumatic brain injury model in the African turquoise killifish (*Nothobranchius furzeri*), a regeneration-competent vertebrate model that evolved to naturally age extremely fast. Stab-wound injury of the aged killifish dorsal telencephalon unveils an impaired and incomplete regeneration response when compared to young individuals. Remarkably, killifish brain regeneration is mainly supported by atypical non-glial progenitors, yet their proliferation capacity appears declined with age. We identified a high inflammatory response and glial scarring to also underlie the hampered generation of new neurons in aged fish. These primary results will pave the way for further research to unravel the factor age in relation to neurorepair, and to improve therapeutic strategies to restore the injured and/or diseased aged mammalian CNS.

**Highlights:**

- Aging impairs neurorepair in the killifish pallium at multiple stages of the regeneration process
- Atypical non-glial progenitors support the production of new neurons in the naive and injured dorsal pallium
- The impaired regeneration capacity of aged killifish is characterized by a reduced reactive proliferation of these progenitors followed by a decreased generation of newborn neurons that in addition, fail to reach the injury site
- Excessive inflammation and glial scarring surface as potential brakes on brain repair in the aged killifish pallium

## Introduction

Age-related neurodegenerative diseases are highly debilitating and incurable pathologies that impinge a high socio-economic burden on our society (El-Hayek *et al*., 2019). They share a progressive degeneration of neurons, which results in loss of brain function and a heterogeneous array of incapacitating symptoms (Dugger *et al*., 2017). Therapeutic strategies for brain restoration consist of compensating for neuronal loss by generating new neurons from the existing stem cell pools that can integrate in the existing circuitry. The capacity for neuroregeneration is naturally limited in the adult mammalian brain (Tanaka *et al*., 2009; Zhao *et al*., 2016). Neural stem cells may start dividing upon injury, but a large part of the newly generated neurons fail to mature, survive and integrate into the existing neural network, thereby restricting full recovery (Arvidsson *et al*., 2002; Thored *et al*., 2006; Kernie *et al*., 2010; Turnley *et al*., 2014; Grade *et al*., 2017). The mammalian neurogenic potential diminishes even further with advancing age, which constitutes one of the main risk factors for neurodegenerative diseases (Galvan *et al*., 2007; Popa-Wagner *et al*., 2011; Hou *et al*., 2019). Studying vertebrate aging in models with high neuroregenerative capacities, such as teleost fish, can therefore help reveal key information to deal with each of these physiological brakes on neuroregeneration and neurorepair (Zhao *et al*., 2016).

Teleost fish share approximately 70% of the coding genes with mammals (Howe *et al*., 2013), but in contrast to mammals, retain the ability to regenerate multiple organs, including fin, heart and the central nervous system (CNS) (Zupanc, 2001; Zupanc *et al*., 2012; Wendler *et al*., 2015; Marques *et al*., 2019; Zambusi *et al*., 2020; Van houcke *et al*., 2021). They have been extensively exploited as gerontology models in the past (Ding *et al*., 2010; Gopalakrishnan *et al*., 2013; Van houcke *et al*., 2015, 2021; Kim *et al*., 2016; Platzer *et al*., 2016). Yet, most teleosts are relatively long-lived (3-5 years) just like mice (Gerhard *et al*., 2002; Miller *et al*., 2002; Gopalakrishnan *et al*., 2013), making investigations about the specific impact of age impractical (Van houcke *et al*., 2021). The African turquoise killifish (*N. furzeri*) however, has surfaced as an ideal vertebrate model for aging studies because of its extremely short lifespan. The short-lived GRZ stain for example, has a median lifespan of 4-6 months depending on housing conditions (Valdesalici *et al*., 2003; Reichwald *et al*., 2015; Valenzano *et al*., 2015; Polačik *et al*., 2016). Killifish naturally live in ephemeral ponds in Africa, which have forced this species to evolve into a short lifespan and rapid aging (Genade *et al*., 2005). Many typical aging hallmarks of mammals are conserved in killifish (Kim *et al*., 2016; Platzer *et al*., 2016; Hu *et al*., 2018; Van houcke *et al*., 2021). Understanding how aged killifish retain or lose their high regenerative capacity upon CNS aging, could thus catalyze the development of therapeutic strategies aiming at inducing successful neuroregeneration in the adult mammalian brain.

The telencephalon or forebrain of fish is a favorable model of study since homologues to the two main neurogenic zones of mammals, the subgranular zone (SGZ) of the hippocampus and the subventricular zone (SVZ) of the lateral ventricles, have been identified in the pallium and subpallium (Adolf *et al*., 2006; Mueller *et al*., 2009, 2011). It was recently discovered that the killifish dorsal pallium holds two classes of progenitors: (1) the commonly known radial glia (RGs) and (2) the non-glial progenitors (NGPs). NGPs are devoid of the typical astroglial/RG markers, such as glutamine synthetase (GS), brain lipid-binding protein (BLBP), glial fibrillary acidic protein (GFAP) and vimentin, and have a morphology that is less branched than RGs (Coolen *et al*., 2020). Early in development, killifish RGs enter a premature quiescent state and proliferation is supported by the NGPs. This is in sharp contrast to the situation in zebrafish, where RGs represent the neurogenic population in the dorsal pallium (Kroehne *et al*., 2011; Rothenaigner *et al*., 2011; Coolen *et al*., 2020). How these two different progenitor classes, with the NGPs appearing unique to the killifish, behave in the neuroregenesis process, and whether their neurogenic capacity is influenced by aging, remains unexplored.

In the present study, we have first set up and validated stab-wound injury as a reliable traumatic brain injury (TBI) model. Next, we have decoded the impact of aging on the regeneration capacity - and on the two neurogenic pools - in the killifish dorsal pallium after stab-wound injury. We find that aged killifish, just like mammals, are not capable to successfully regenerate; they develop an excessive inflammatory reaction and glial scarring, show diminished injury-induced neurogenesis, and the vast majority of newborn neurons fail to reach the injury site. Remarkably, we show neuroregeneration to be supported by the NGPs in both young adult and aged killifish, and not the RGs that typically constitute neuroregenesis in other teleosts. Taken together, aged killifish appear to mimic the impaired regeneration capacity also seen in adult and/or aged mammals, instead of displaying the high regenerative capacities seen in young adult teleost fish, including killfish. In summary, we propose the killifish aging nervous system as a valuable model to create knowledge about the identity and mode of action of drivers and brakes of neuroregenerative properties.This may eventually elucidate how to boost the repair capacity in an aging context in the diseased/injured mammalian brain.

## Methods

### RESOURCE AVAILABILITY

#### Lead contact and Materials availability

Further information and requests for resources (killifish) should be directed to and will be fulfilled by the Lead Contact, Lutgarde Arckens.

#### Data and code availability

The present study has no unique datasets or code.

## EXPERIMENTAL MODEL AND SUBJECT DETAILS

### Fish strain and housing

All experiments were performed on adult (6 week- and 18 week-old) female African turquoise killifish (*Nothobranchius furzeri*), inbred strain GRZ-AD, which were kindly provided by Prof. Dr. L. Brendonck and Dr. T. Pinceel and originate from the Biology of Ageing, Leibniz Institute for Age Research - Fritz Lipmann Institute, Jena, Germany. Breeding pairs were housed in 8 L aquaria and experimental fish were kept in 3,5 L aquaria in a ZebTEC Multi-Linking Housing System (Tecniplast). One male was housed with three females under standardized conditions; temperature 28 °C, pH 7, conductivity 600 μs, 12h/12h light/dark cycle, and fed twice a day with (*Artemia salina*, Ocean Nutrition) and mosquito larvae (*Chironomidae*, Ocean Nutrition). Breeding pairs were given sandboxes for spawning. Fertilized eggs were collected once a week and washed with methylene blue solution (0,0001% in autoclaved system water, Sigma-Aldrich, 03978) for five minutes. Next, eggs were bleached twice for five minutes in 1% Hydrogen Peroxide (H_2_O_2_, Chem-Lab, CL00.2308.5000) diluted in autoclaved system water. Then, eggs were again washed four times for five minutes with methylene blue solution. Eggs were stored on moist Jiffy-7C coco substrate plates (Jiffy Products International AS, Norway) at 28°C with a 12h/12h light/dark cycle for three weeks in a custom-made incubator. Embryos that reached the ‘Golden Eye stage’ (Polačik *et al*., 2016) were hatched in a small volume of ice-cold Humic acid solution (Sigma-Aldrich, 53680, 1g/L in system water) with continuous oxygenation. Larvae were raised at 26°C and half of the water was changed daily for one week. Hereafter, larvae were transferred to 3,5 L aquaria and fed daily with (*Artemia salina*, Ocean Nutrition) until 3 weeks post hatching. All experiments were approved by the KU Leuven ethical committee in accordance with the European Communities Council Directive of 22 September 2010 (2010/63/EU) and the Belgian legislation (KB of 29 May 2013).

## Method details

### Stab-wound injury

Fish were first sedated in 0,03% buffered tricaine (MS-222, Sigma-Aldrich, CAS: 886-86-2), diluted in system water, and placed in a cold moist sponge. Scales, skin and fat tissue was removed above the right telencephalic hemisphere to visualize the skull. Next, a custom Hamilton 33-Gauge needle was pushed through the skull into the dorsal pallium of the right hemisphere of the telencephalon, causing a brain lesion of approximately a depth of 500 μm (Figure S1). The telencephalon lies in between the eyes of the killifish, which were used as landmarks. The needle was dipped in Vybrant DiD cell-labeling solution (C_67_H_103_ClN_2_O_3_S, Thermo Fisher Scientific, V22887) to easily find the place of entrance and needle track on sections (Figure S1). Afterwards, the fish were placed in fresh system water to recover.

### BrdU labeling

To label dividing cells and their progeny, fish were placed in 5-Bromo-2’-deoxyuridine (BrdU, Sigma-Aldrich, B5002-5G; CAS: 59-14-3) water (7,5 mM in system water) for 16 hours between one and two days post injury. After the pulse, fish were placed in fresh system water for a chase period of 21 days.

### Tissue fixation and processing

Fish were euthanized in 0.1% buffered tricaine (MS-222, Sigma-Aldrich, diluted in system water) and perfused via the heart with PBS and 4% paraformaldehyde (PFA, Sigma-Aldrich, 8.18715, CAS 30525-89-4, diluted in PBS). Brains were extracted and fixed for 12 hours in 4% PFA at 4 °C. Afterwards, brains were washed three times with PBS and embedded in 30% sucrose, 1,25% agarose in PBS. Coronal sections of 10 µm were made on a CM3050s cryostat (Leica) and collected on SuperFrost Plus Adhesion slides (Thermo Fisher Scientific, 10149870). Sections were stored at -20 °C until immunohistochemistry (IHC) or Cresyl Violet staining.

### Cresyl Violet histological staining

Cryostat sections were dried for 30 minutes at 37°C to improve adhesion to the glass slides and washed in *aqua destillata* (AD). Next, the sections were immersed in Cresyl Violet solution (1% in AD, Fluka Chemicals, Sigma-Aldrich) for five minutes. The sections were rinsed in 200 mL AD with five drops of Acetic Acid (Glacial, 100%) for 30 seconds. Hereafter, the sections were dehydrated in 100% ethanol and 100% xylol series. Sections were covered with DePeX and cover slip and dried overnight.

### Immunohistochemistry (IHC)

Sections were dried for half an hour at 37°C to improve adhesion to the glass slides and washed in AD and TBS (0,1% Triton-X-100 in PBS). Heat-mediated antigen retrieval was used to break cross-links between the proteins. The slices were boiled in the microwave in 1X citrate buffer (2,1 g citric acid and 500 µL Tween 20 in 1L PBS, pH 6) for five minutes at 100% and two times five minutes at 80%. Afterwards, the slices were cooled down for 20 minutes and washed three times for five minutes with TBS. For BrdU IHC, sections were pretreated with 2N HCl at 37°C for 30 minutes to break the DNA and washed with 0,1M sodium borate (in AD) to neutralize HCl. Sections were blocked for one hour at room temperature with 20% normal goat serum (Sigma-Aldrich, S26) in Tris-NaCl blocking buffer (TNB). For IHC stainings involving the primary antibody Goat anti-BLBP (Abcam, ab110099), blocking was performed with normal donkey serum (Sigma-Aldrich, S30). Sections were stained over night with primary antibodies diluted in TNB at room temperature, with the exception of the anti-HuC/D antibody, which was incubated at 4°C for 48 hours in Pierce Immunostain Enhancer (Thermo Fisher Scientific, 46644). Pierce Enhancer was also used in triple IHC stainings and when the anti-L-plastin primary antibody (GeneTex, GTX124420) was involved. The primary antibodies used were rabbit anti-SOX2 (1:1000, Sigma-Aldrich, SAB2701800), mouse anti-HuC/D (1/200, Thermo Fisher Scientific, A-21271), mouse anti-PCNA (1:500, Abcam, ab29), Goat anti-BLBP (1:1000, Abcam, ab110099), rat anti-BrdU (1:1000, Abcam, ab6326), rabbit anti-L-plastin (1:400, GeneTex, GTX124420) and mouse anti-GS (1:1000, Abcam, ab64613). Sections were washed with TBS three times for five minutes. Secondary antibodies were stained at room temperature in TNB for two hours (Alexa-594, Alexa-488, Alexa-305, 1:300, Thermo Fisher Scientific). In case of the anti-L-plastin IHC staining, a long amplification was used, in which the secondary antibody is coupled to biotin (Goat anti-Rabbit-biotin, 1:300 in TNB, Agilent Dako, E043201-8) and incubated for 45 minutes. After washing three times for five minutes with TBS, Streptavidin-Cy5 (1/500 in TNB, Thermo Fisher Scientific, SA1011) was added to the sections for two hours. For cell death detection, the TUNEL assay (In Situ Cell Death Detection Kit, Fluorescein, Sigma-Aldrich, 11684795910) was used, following the manufacturer’s instructions. Cell nuclei were stained with 4’,6-diamidino-2-fenylindool (DAPI, 1:1000 in PBS, Thermo Fisher Scientific). Last, sections were covered with Mowiol solution and a cover glass slide.

### Microscopy

Sections were scanned for DiD positivity using Texas Red light with a confocal microscope (FV1000, Olympus), to locate the injury site, even after neuroregeneration was completed. For the quantification of IHC stainings and Cresyl Violet staining, a Zeiss (‘Axio Imager Z1’) fluorescence microscope was used to photograph three sections per animal. For fluorescence pictures the AxioCam MR R3 camera was used. For Bright field pictures, the Mrc5 color camera was used. 20X Tile scans were taken and stitching was performed using the ZEN software (ZEN Pro 2012, Carl Zeiss). For greater detail, 63X magnification pictures were taken using immersion oil. Channels were equally intensified for all conditions using Adobe Photoshop SC5 (Adobe Systems) for publication. Figure configurations were made with Adobe Illustrator (Adobe Systems).

## QUANTIFICATION AND STATISTICAL ANALYSIS

Immunopositive cells were counted (Figure S4) and the injury area surface (mm^2^) was measured (Figure S2) on three sections adjacent to the lesion site and compared with corresponding sections of naive animals using respectively the cell counter plugin and polygon tool in Image J (Fiji). An average of three sections was taken per animal for statistical analysis using GraphPad Prism (version 8.2.1).

Data were analyzed by two independent observers and first tested for Gaussian normality and homoscedasticity. If these assumptions were met, we used a parametric unpaired *t*-test (two conditions) or one-way ANOVA (> two conditions), followed by Dunnett’s multiple comparisons test to compare injured fish to naive fish for each age separately. If these assumptions were not met, we used the non-parametric Mann Whitney test (two conditions) or Kruskal-Wallis test (> two conditions), followed by Dunn’s multiple comparisons test. Two-way ANOVA was used to compare young and aged fish at each time point, followed by Sidak’s multiple comparisons test. n represents the number of animals in each condition. All values are mean ± standard error of the mean (SEM). Means were statistically significantly different when p≤ 0,05.

## Results

### A stab-wound injury model to study neuroregeneration in the killifish pallium

Before studying neuroregeneration in the young and aged dorsal pallium of killifish, we first introduced an easy-to-use and reproducible brain injury model. Stab-wound injuries have been extensively characterized in the zebrafish telencephalon and effectively induce brain regeneration (Ayari *et al*., 2010; Kroehne *et al*., 2011; März *et al*., 2011; Baumgart *et al*., 2012; Kishimoto *et al*., 2012; Kyritsis *et al*., 2012; Barbosa *et al*., 2015). We chose to optimize a similar injury model in the young adult (6 week-old) and aged (18 week-old) killifish telencephalon (Figure S1A-B). Approaching the dorsal telencephalon through the nostrils was not possible without damaging the large eyes, since the nostrils of the killifish are positioned more lateral compared to zebrafish. Instead, the medial part of the right telencephalic hemisphere was targeted from above, and a 33-gauge Hamilton needle was pushed through the skull, approximately 500 µm in depth (Figure S1C). The medial telencephalon lies in between the eyes, which were used as visual landmarks. Skin and fat tissue cover the skull of the killifish and were removed to visualize the skull of the fish. The procedure required an experienced hand and a training period. Just prior to wounding, the needle was dipped in Vybrant DiD Cell-Labeling Solution (Thermo Fischer) in order to permanently label the cells close to the injury site. This DiD dye approach enabled reconstruction of the original injury site, even if complete recovery had taken place, and the wounded area could no longer be distinguished from the surrounding tissue by (immuno)histology (Figure S1D,E).

### Aging impairs tissue recovery after stab-wound injury

Standard histology revealed a packed blood clot that filled the injury site in the brain of young adult and aged killifish immediately upon injury (Figure 1A,B). In ensuing weeks, the parenchyma of the young adult telencephalon was able to structurally regenerate in a seamless fashion, showing a normal distribution of cells even at the DiD-positive injury site, 23 to 30 days post injury (dpi) (Figure 1C, see also Figure S1F). In aged killifish on the contrary, the parenchyma showed a malformation, recognizable by swollen and irregularly-shaped cells and blood vessels at 30 dpi. The malformation was reminiscent of a mammalian glial scar (Figure 1D, see also Figure S1G). We tested if the scar tissue, only observed in aged killifish, was of glial nature. As the teleost telencephalon is devoid of parenchymal astrocytes (Grupp *et al*., 2010), we probed for the presence of microglia and the long bushy fiber of RGs (GS^+^) that span the entire parenchyma. At 23 dpi a cluster of L-plastin^+^ microglia/macrophages was present at the injury site, co-localising with the scar tissue and the DiD labeled cells in aged killifish. Even at 30 dpi, the microglia/macrophage cluster was still present in aged injured killifish (Figure S1F-I) indicating that the scar is likely permanent. We did not observe RG cell bodies at the scar tissue. However, the GS^+^ fibers of the RG were surrounding the glial scar, indicating that RGs were still involved in glial scarring via the use of their bushy fiber.

**Figure 1:**
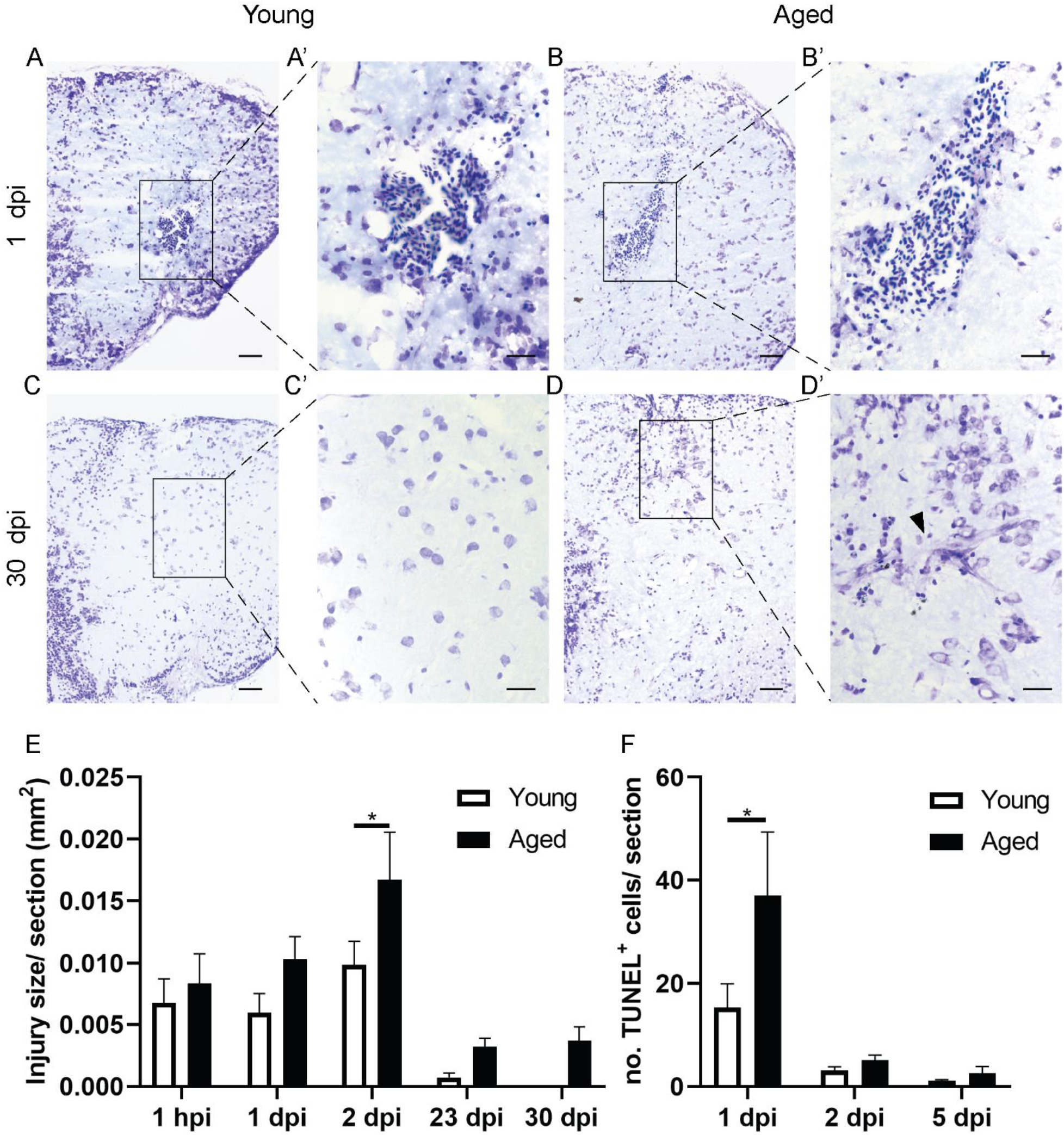
Aging impairs tissue recovery after brain injury. **(A-D)** Cresyl violet stainings of coronal sections illustrate the injury site and tissue recovery after brain injury in the telencephalon of young killifish (A,C) and of aged killifish (B, D). At 30 dpi, the aged telencephalon still shows a tissue scar in the parenchyma (arrowhead in D’), while the young telencephalon has no visible injury anymore. Of note, cells near the tissue scar appear swollen (D’). Scale bar in A-D: 50 µm. Scale bar in A’-D’: 20 µm. **(E)** Quantification of the injury surface area (in mm^2^) measured at 1 hpi and 1, 23, 30 dpi, in young adult and aged killifish. At 1 hpi the injury size is similar between young adult and aged killifish, towards 2 dpi the injury is significantly enlarged in aged fish. In addition, the injury is still visible at 23 and 30 dpi, time points at which young adult fish demonstrate extensive structural recovery. **(F)** Countings of the absolute number of TUNEL^+^ apoptotic cells detected in the injured telencephalon at 1, 2 and 5 dpi reveal that apoptosis is significantly larger in aged fish at 1dpi. *p≤0,05; Two-way ANOVA, followed by Sidak’s multiple comparisons test. Values are mean ± SEM; n≥4. hpi: hours post injury, dpi: days post injury. Pictures of coronal brain sections in (A, B,C,D) are also shown in Figure S2 to illustrate how the injury surface area was measured, and in Figure S1 to visualize the DiD crystals and glial scarring at the site of injury.

In conclusion, stab-wound injury could effectively be applied to compare the regeneration process between young and aged killifish brains. Aged killifish brain recovered incompletely and showed signs of permanent glial scarring.

### A high inflammatory response exacerbates tissue recovery in aged killifish

To be certain that the stab-wound injury created a comparable injury in young and aged animals, we studied the temporal dynamics of the surface size of the injury using (immuno)histology (Figure S2). One hour after injury, the size of the wound was comparable between young adult and aged killifish. By 2 dpi, the injury size was enlarged in aged killifish, possibly due to differences in cell death, arrival of inflammatory cells and edema (Figure 1E). By counting the number of TUNEL^+^ apoptotic cells we discovered that the aged, injured brain contained more TUNEL^+^ apoptotic cells than young adult fish already at 1 dpi (37±12,33 versus 15,33±4,659; P=0,0128). It thus seems that in an aged injured environment, cells were more vulnerable to secondary damage, which might be linked to an inflammatory state. Led by these results, we stained for the microglia/macrophage marker (pan-leukocyte marker) L-plastin at 1 and 2 dpi. As expected, stab-wound injury induced an acute inflammatory response reflected in an increased number of microglia/macrophages at 1 and 2 dpi in injured fish compared to naive fish (uninjured controls) (Figure 2). In addition, we detected more microglia/macrophages in aged, injured killifish compared to young adult, injured killifish, indicating a higher inflammatory response in aged killifish (Figure 2). This may explain the higher number of apoptotic cells seen at 1 dpi, as well as why the size of the injury site increases towards 2 dpi in aged killifish. In addition, a fraction of these microglia/macrophages were PCNA^+^ (Figure S3), and thus proliferating, which is in accordance with reports on injury impact in zebrafish (Kroehne *et al*., 2011; Baumgart *et al*., 2012).

**Figure 2:**
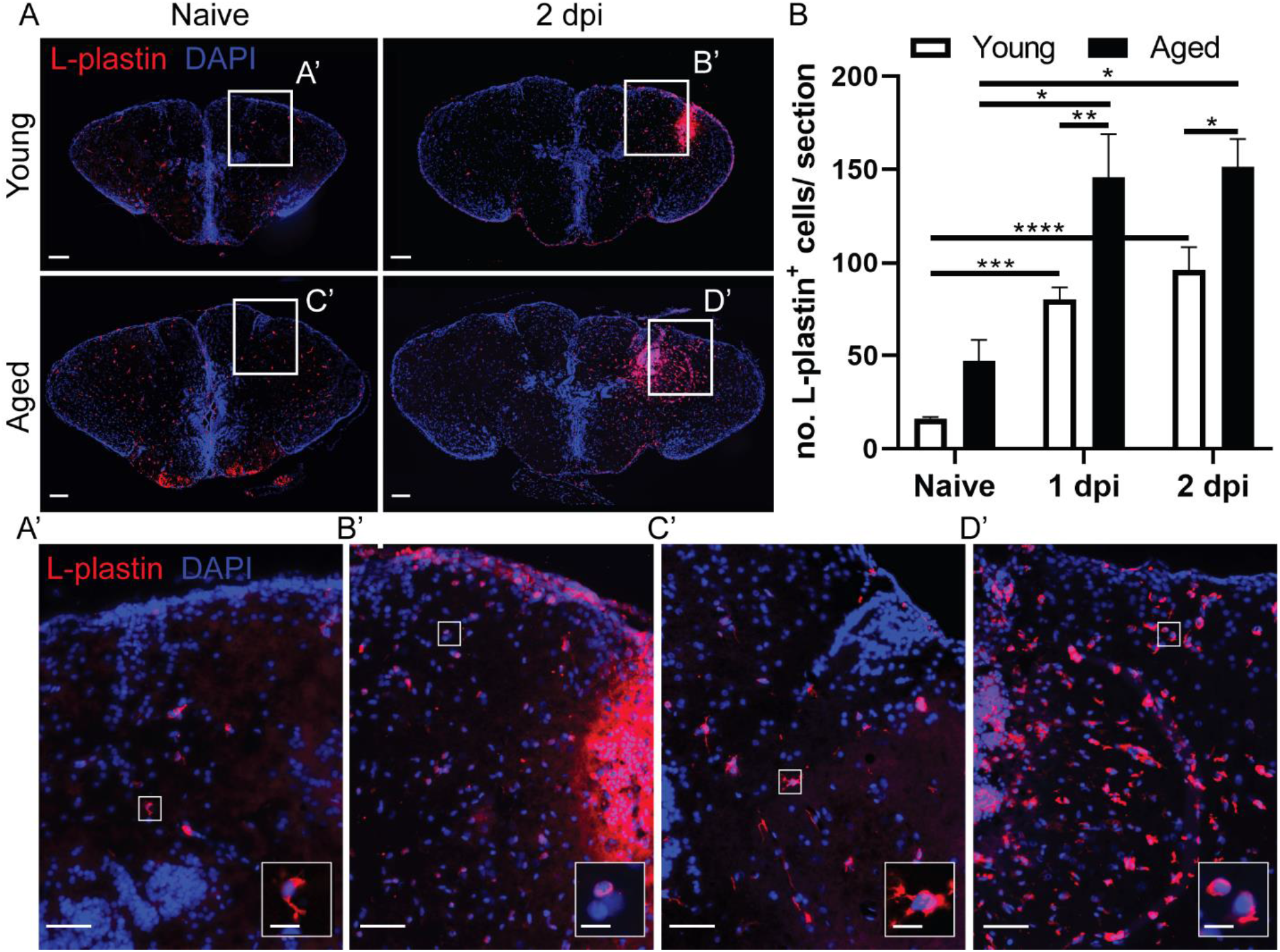
The injury-induced inflammatory reaction is larger in the aged brain. **(A)** Staining for L-plastin (red) with DAPI (blue) on coronal brain sections of young adult and aged killifish in naive conditions and at 2 dpi. Boxed areas are magnified in (A’-D’). A more pronounced increase in L-plastin^+^ inflammatory cells is observed for aged than young adult injured fish. (A’-D’) Higher magnification of individual L-plastin^+^ microglia/macrophages are presented in the right bottom corner. Scale bars in (A) and (A’-D’): 100 µm. Scale bars of boxed areas in A’-D’: 10 µm. **(B)** Absolute number of L-plastin^+^ microglia/macrophages in young adult and aged telencephali in naive conditions and at 2 dpi. Both young adult and aged killifish show a significant increase in inflammatory cells early after brain injury, but the number of microglia/macrophages is significantly higher in aged fish at 1 and 2 dpi when compared to young adult individuals. *p≤0,05, **p≤0,01, ***p≤0,001, ****p≤0,0001; One-way ANOVA is used to compare naive fish to injured fish. Young: parametric one-way ANOVA, followed by Dunnett’s multiple comparisons test. Aged: non-parametric Kruskal-Wallis test, followed by Dunn’s multiple comparisons test. Two-way ANOVA is used to compare young and aged fish, followed by Sidak’s multiple comparisons test. Values are mean ± SEM; n≥5. dpi: days post injury.

### Reactive proliferation is declined and delayed in aged injured killifish brains

For brain repair to be successful, new neurons should be generated from the available neuronal progenitor cell (NPC) pool. We counted the number of activated NPCs (SOX2^+^ PCNA^+^) and all NPCs (SOX2^+^) in the dorsal ventricular zone (VZ) (Figure S4) in function of age and time post injury.

The percentage of dividing NPCs in the progenitor pool of aged naive fish was cleary lower than in young fish (8,120%±1,048 versus 20,06%±3,486; P=0,0047, Figure 3), and this difference persisted upon injury. In young killifish, the injury-induced proliferation of NPCs was significantly higher than in aged killifish (at 2 dpi: 34,94%±3,824 versus 17,75%±2,629 repectively; P<0,0001, Figure 3). The increase occured at 1 and 2 dpi, and declined back to normal levels from 5 dpi onward. In the aged killifish the percentage was significantly higher at 2 dpi compared to naive aged fish, but it did not even reach the baseline levels of the young adult killifish. These result show that the capacity for NPC proliferation upon injury diminishes steeply with age, but that aged killifish still retain some capacity for NPC reactive proliferation.

**Figure 3:**
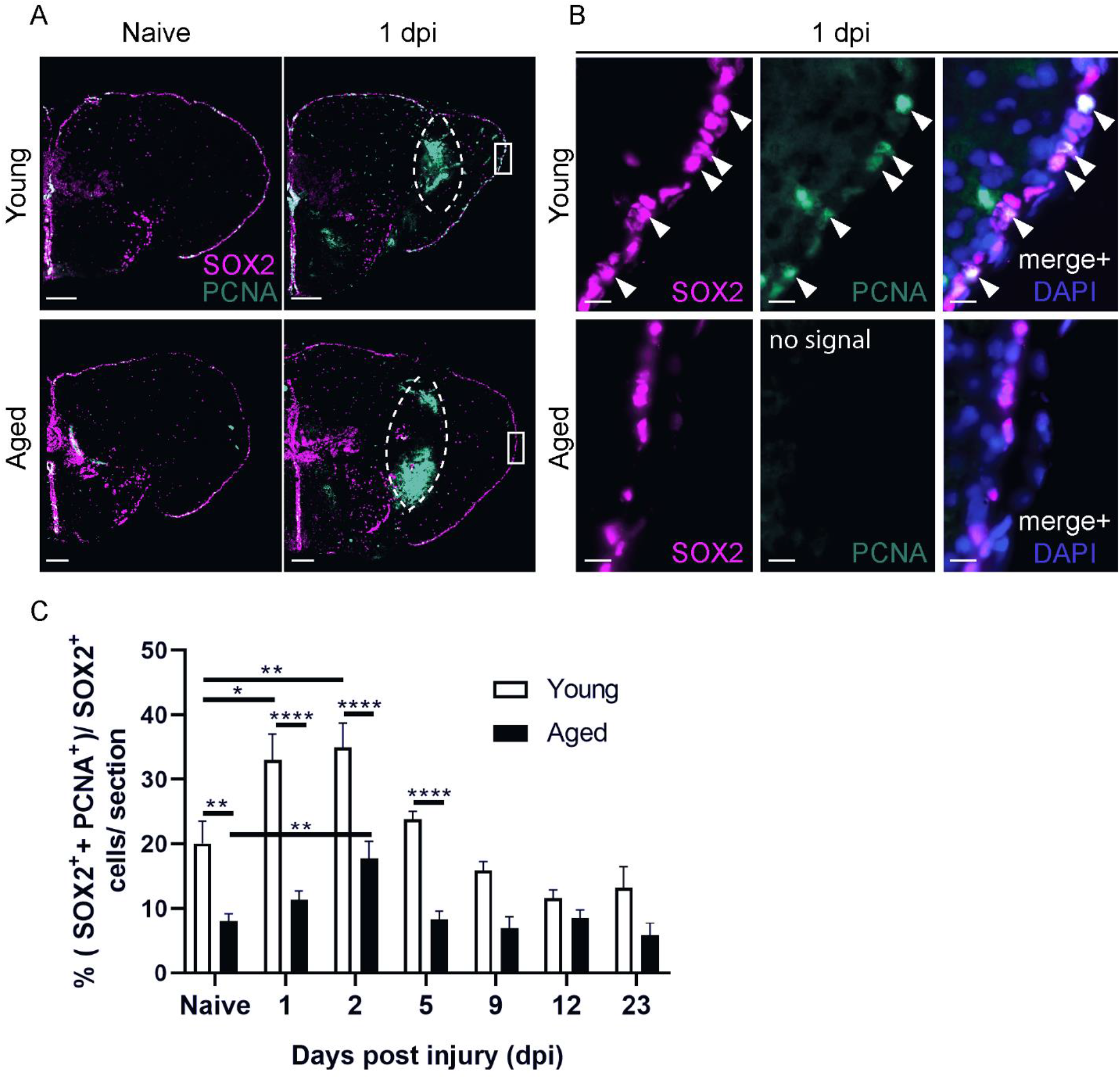
Aging diminishes neural progenitor cell proliferation in the ventricular zone of the killifish telencephalon, in naive conditions and in response to injury. **(A, B)** Double staining for SOX2 (magenta) and PCNA (green) with DAPI (blue) on coronal sections of young and aged killifish in naive conditions and at 1 dpi. The dashed lines encircle the site of injury filled with blood cells that autofluoresce in the green channel. **(B)** Magnifications of the boxed areas in A: Young adult killifish have a higher percentage of proliferating progenitor cells (SOX2^+^ PCNA^+^ cells, arrowheads) in the VZ compared to aged fish. Scale bars in A: 100 µm. Scale bars in B: 10 µm. **(C)** Proportion of double-positive SOX2^+^ PCNA^+^ dividing progenitor cells among SOX2^+^ cells (all progenitor cells) in young adult and aged telencephali in naive conditions and at 1, 2, 5, 9, 12 and 23 dpi. At all time points investigated, young adult fish have a higher capacity for progenitor proliferation than aged fish. Interestingly, for both ages progenitor proliferation peaks at 2 dpi. *p≤0,05, **p≤0,01, ****p≤0,0001; One-way ANOVA is used to compare naive fish to injured fish, followed by Dunnett’s multiple comparisons test. Two-way ANOVA is used to compare young and aged fish at each time point, followed by Sidak’s multiple comparisons test. Values are mean ± SEM; n≥5, except for aged, 9 dpi: n=4. VZ: ventricular zone, dpi: days post injury.

### Reactive proliferation is mainly supported by NGPs and not RGs

The killifish dorsal telencephalon holds two classes of progenitors: NGPs and RGs (Coolen *et al*., 2020). We therefore investigated which progenitor type supported reactive proliferation in our injury model at 2 dpi, the time point at which reactive proliferation peaks independent of age (Figure 3). By immunohistochemistry for SOX2, PCNA and BLBP, we could delineate the dividing RGs (SOX2^+^ PCNA^+^ BLBP^+^) from the dividing NGPs (SOX2^+^ PCNA^+^ BLBP^-^). Strikingly, we rarely observed dividing RGs (approximately 3% of all NPCs) (Figure 4), although RG fibers did present themselves with a swollen morphology after injury (Figure 4B’). Instead, we clearly observed many dividing NGPs in the naive and injured dorsal VZ of young and aged killifish, showing that NGPs are the most prominent cell type supporting adult neurogenesis in killifish (Figure 4). The percentage of dividing NGPs among all NPCs was significantly lower in aged killifish compared to young adult fish (Figure 4), both in naive animals (14,26%±1,328 versus 28,04%±1;569; P=0,0005) and injured animals (22,5%±2,383 versus 43%±2,543; P<0,0001), demonstrating that adult neuro(re)genesis declined upon aging.

**Figure 4:**
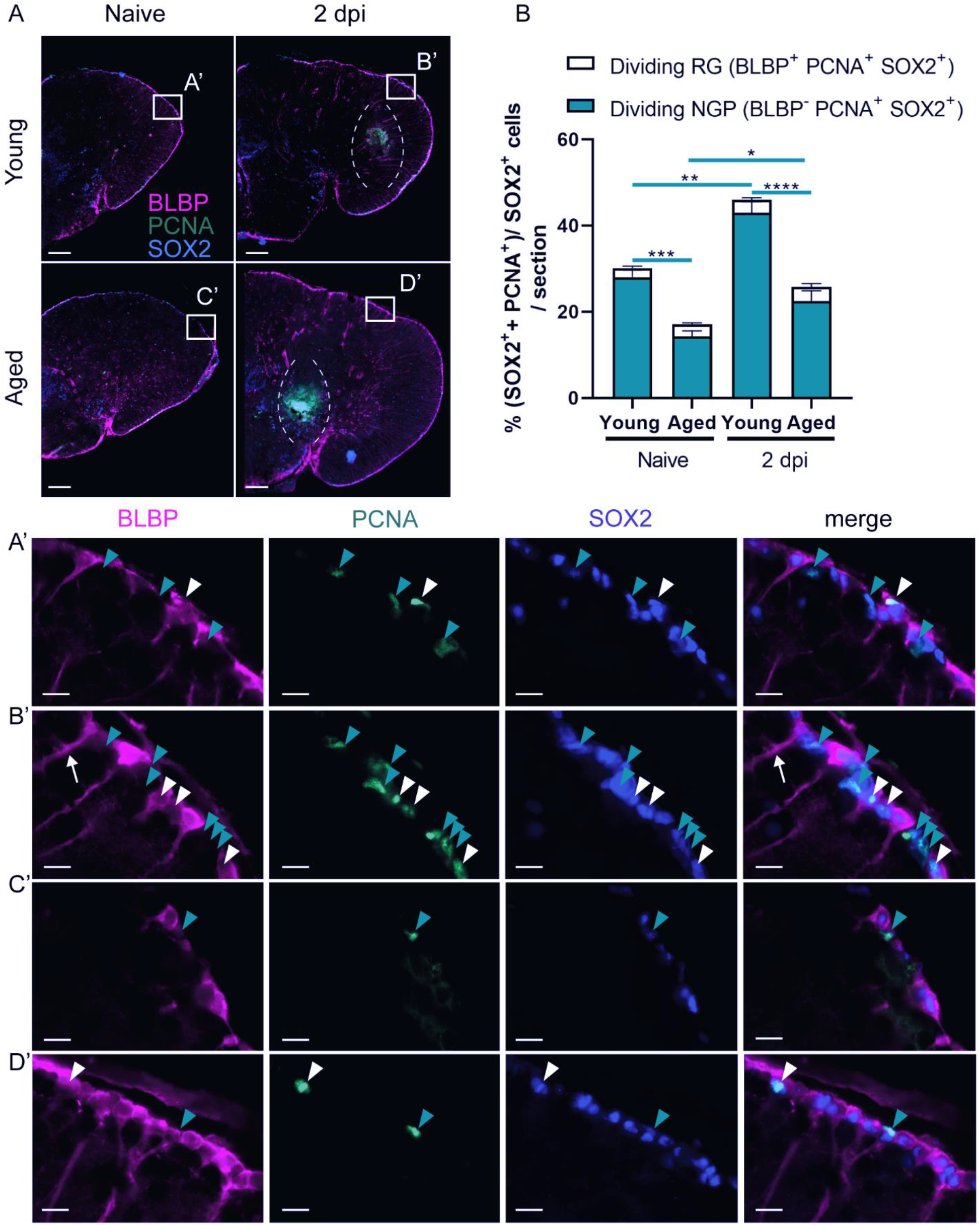
Aging reduces the proportion of dividing specialized NGPs, but not dividing common RGs. **(A)** Triple staining for BLBP (magenta), PCNA (green) and SOX2 (blue) on coronal sections of young adult and aged killifish in naive conditions and at 2 dpi. The dashed lines encircle the site of injury filled with blood cells that autofluoresce in the green channel. Boxed areas are magnified in (A’-D’). White arrowheads depict triple positive BLBP^+^ SOX2^+^ PCNA^+^ dividing RGs, while turquoise arrowheads mark double positive BLBP^-^ SOX2^+^ PCNA^+^ dividing NGPs. (C’) Thicker RG fibers are noticed in young, injured fish (white arrow in B’), indicative for glial activation. Young adult killifish clearly have more dividing NGPs compared to aged fish. In addition, dividing RGs are only observed in small amounts in the VZ. Scale bars in (A): 100 µm. Scale bars in (A’-D’): 10 µm. **(B)** Proportion of BLBP^+^ SOX2^+^ PCNA^+^ dividing RG and BLBP^-^ SOX2^+^ PCNA^+^ dividing NGPs over all SOX2^+^ cells (all progenitor cells) in the VZ of young and aged telencephali in naive conditions and at 2 dpi. Reactive progenitor proliferation is mainly supported by specialized NGPs and is highest in young adult killifish. Aged killifish have significantly lower percentages of dividing NGPs in both naive and injured conditions. *p≤0,05, **p≤0,01, ***p≤0,001, ****p≤0,0001; Unpaired t-test is used to compare naive fish to injured fish. Two-way ANOVA is used to compare young and aged fish, followed by Sidak’s multiple comparisons test. Values are mean ± SEM; n≥5. RG: radial glia, NGP: non-glial progenitor, VZ: ventricular zone, dpi: days post injury.

Unlike in zebrafish, we provide evidence that injury-induced reactive proliferation of progenitors is mainly supported by the NGPs in the killifish pallium. Of all dividing NPCs present in the young injured adult VZ at 2 dpi, 93,49%±0,688 could be assigned to NGPs, while only 6,509%±0,688 were of RG type (Table S1). This is in sharp contrast to young adult injured zebrafish where 86,5%±3,1% (n=4) of dividing cells represented RGs at 3 dpi (Kroehne *et al*., 2011). In the aged injured killifish 87,72%±1,837 of all dividing NPCs were NGPs, while 12,28%±1,837 were RG (Table S1). Aged injured killifish thus showed a higher percentage of dividing RG among all dividing cells compared to young adult injured killifish. A similar observation was made for naive conditions (Table S1). It appears that aged killifish try to compensate their reduced capacities by activating the RG pool.

Taken together, killifish NGPs represent the most potent progenitor type driving the proliferative response to injury, but their number declines significantly with age when RGs appear more activated, albeit still low in repect to NGPs.

### A declined number of newborn neurons reach the injury site in aged killifish

To elucidate if reactive NGPs lead to the production of newborn neurons that can migrate to the injury site for replacement of the lost cell types, we designed a BrdU pulse chase experiment. In between 1 and 2 dpi, when reactive proliferation is most intense (Figure 3), young and aged injured and age-matched control (AMC) killifish where subjected to 16 hours of BrdU water, labeling all cells passing through S-phase (Figure 5A). After a 21-day chase period, a time window matching maximal recovery in young adult fish, we visualized the progeny of these cells via immunostaining for BrdU and HuCD. HuCD is a pan-neuronal marker, expressed in both early (immature) and late (mature) neurons (Kim *et al*., 1996; Kroehne *et al*., 2011). As such, BrdU^+^ HuCD^+^ neurons, that represent the progeny of the dividing NGPs, were visualized at 23 dpi (Figure 5B).

**Figure 5:**
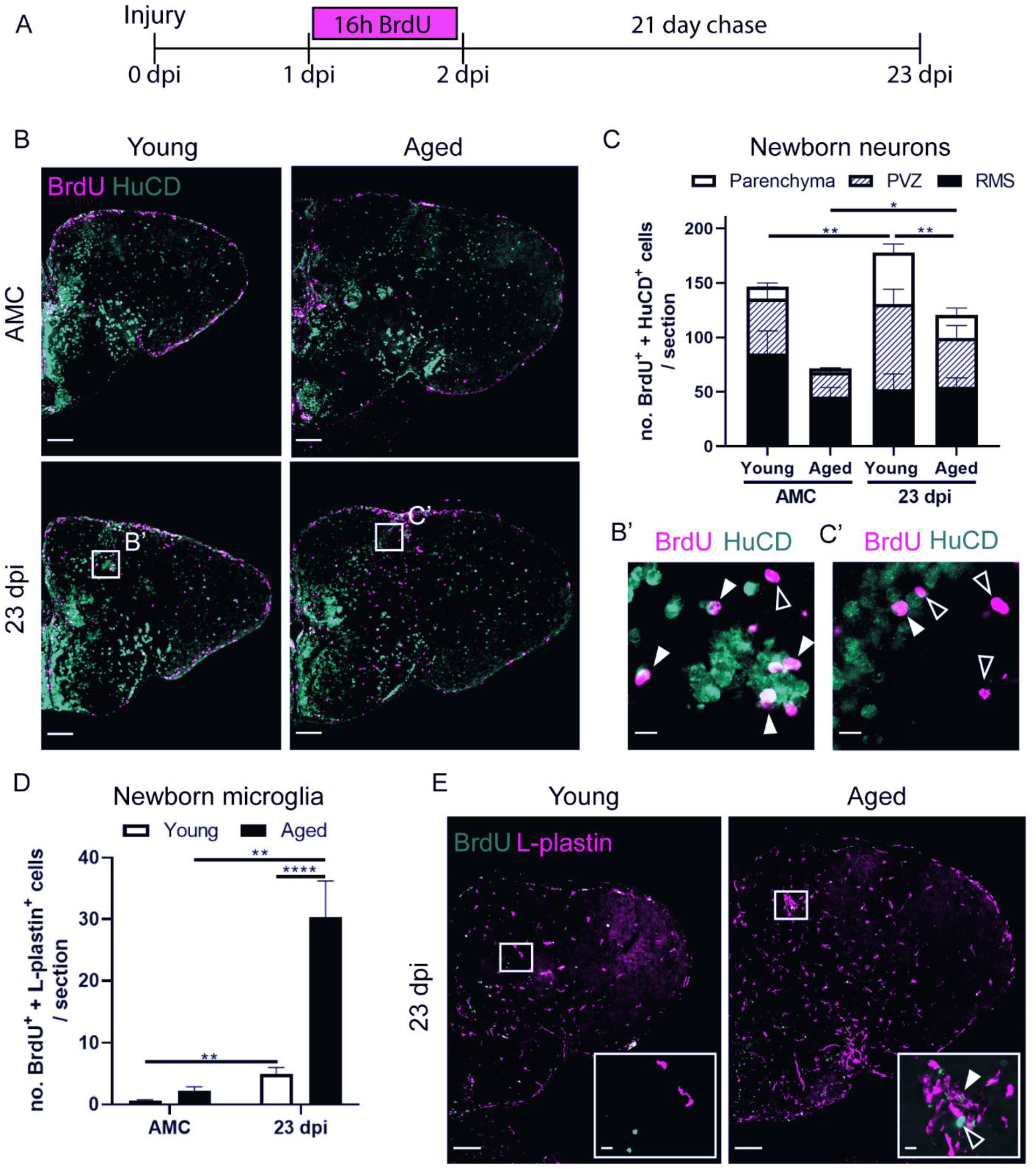
Aged killifish generate less newborn neurons but more newborn microglia/macrophages in the parenchyma of the telencephalon compared to young ault killifish. **(A)** Experimental set up: Day 0: time of a stab-wound injury; From 1 to 2 dpi: injured fish and AMCs are placed in BrdU water for 16 hours: BrdU will be incorporated in the DNA of dividing cells and passed on to the progeny upon each cell division. 23 dpi: regeneration is completed in young fish, all brain samples of young and aged, AMC and injury conditions are collected for cell analysis (as illustrated in B-E). **(B)** Double staining for BrdU (magenta) and HuCD (green). Boxed areas are magnified in (B’,C’). White closed arrowheads depict double positive BrdU^+^ HuCD^+^ newborn neurons; open arrowheads indicate newborn single positive BrdU^+^ cells of unknown cell type. While the BrdU signal overlaps with HuCD (neuronal marker) in young adult killifish, representing newborn neurons, aged killifish mostly have BrdU positive cells lying next to HuCD^+^ neurons, suggesting these cells are another cell type. Scale bars in (B): 100 µm. Scale bars in (B’,C’): 10 µm. **(C)** Number of BrdU^+^ HuCD^+^ newborn neurons near the RMS of the subpallium, in the PVZ of the dorsal pallium, and in the parenchyma. In the parenchyma, significantly lower numbers of newborn neurons are present in aged injured fish compared to young adult injured fish. **(D)** Number of BrdU^+^ L-plastin^+^ newborn microglia/macrophages in the parenchyma. Aged injured fish generate large numbers of newborn microglia/macrophages by 23 dpi. *p≤0,05, **p≤0,01, ****p≤0,0001; Unpaired t-test or non-parametric Mann Whitney test is used to compare AMC to 23 dpi fish. Two-way ANOVA is used to compare young and aged fish, followed by Sidak’s multiple comparisons test. Values are mean ± SEM; n≥5. **(E)** Double staining for BrdU (green) and L-plastin (magenta). Boxed areas are magnified in the right corner of each panel. A cluster of L-plastin^+^ microglia/macrophages is visible in the parenchyma of aged injured fish, but not in young injured fish, at 23 dpi. White closed arrowhead depicts a BrdU^+^ L-plastin^+^ newborn microglia/macrophages, inside this cluster. The open arrowhead points to a green autofluorescent blood cell recognizable by its oval shape and visible nucleus. Scale bars in (E): 100 µm. Scale bars of boxed areas: 10 µm. 16h: 16 hours, AMC: age-matched control, RMS: rostral migratory stream, PVZ: periventricular zone, dpi: days post injury.

Our data reveal impaired production and migration of newborn neurons in aged killifish at 23 dpi. In naive fish, we could hardly detect newborn neurons that had migrated into the parenchyma (10,94±3,303 for young adult and 3,571±0,566 for aged fish) (Figure 5C). Many newborn neurons in the periventricular zone (PVZ) of the dorsal pallium and near the rostral migratory stream (RMS) (Figure S4), migrate only 1-2 cell diameters away from the ventricular stem cell zones, which is typical for constitutive neurogenesis in teleosts (Adolf *et al*., 2006; Grandel *et al*., 2006; Rothenaigner *et al*., 2011). Injured killifish on the contrary, generated more newborn neurons that migrated into the injured parenchyma. This number was significantly higher in young adult killifish than in aged killifish (46,97±8,017 versus 20,80±6,540; P=0,005) (Figure 5C).

Most likely the impaired replenishment of newborn neurons in aged injured animals is caused by a declined production of neurons, yet also the failure of these neurons to migrate towards the injury site through the aged non-permissive environment. Indeed, we confirmed that aged injured killifish produce large numbers of newborn microglia/macrophages (BrdU^+^ L-plastin^+^ cells) by 23 dpi (aged killifish: 30,33±5,88 versus young adult killifish: 4,933±1,087; P<0,0001), what most likely contributes to an inflammatory non-permissive environment (Figure 5D,E).

### Aging hampers the replenishment of progenitors in the VZ after injury

Stab-wound injury disrupts the parenchyma of the dorsal telencephalon, but also the VZ. Hence, the dorsal VZ needs to be replenished with newly generated NGPs and RGs. We applied BrdU pulse chase labeling to investigate if dividing progenitors, labeled between 1 and 2 dpi, give rise to new progenitors by 23 dpi, the time point where regeneration is complete in young adult fish (Figure 6A). We zoomed in on the two major progenitor classes; (1) newly generated RGs (BrdU^+^, BLBP^+^) and (2) newly generated dividing NGPs (BrdU^+^, PCNA^+^, BLBP^-^). The BrdU signal in highly proliferative progenitors, which are NGPs, will be diluted with each cell division until the moment when the BrdU signal gets lost. This assay thus rather describes the progeny of low proliferative progenitors, that is the RGs in killifish. Indeed, in all conditions we found a larger number of newly generated RGs than NGPs. RGs, labeled between 1 and 2 dpi, thus seem to generate new RGs via gliogenic divisions. Whether RGs also gave rise to NGPs or vice versa remains elusive and an interesting research question for future studies. Of note, we also detected a low number of BrdU^+^ cells in the VZ that were PCNA^-^ and BLBP^-^, which we thus could not identify as RG or NGP. We however realize that these cells could possibly represent a quiescent progenitor type or newborn neuroblasts, lying closely to the VZ.

**Figure 6:**
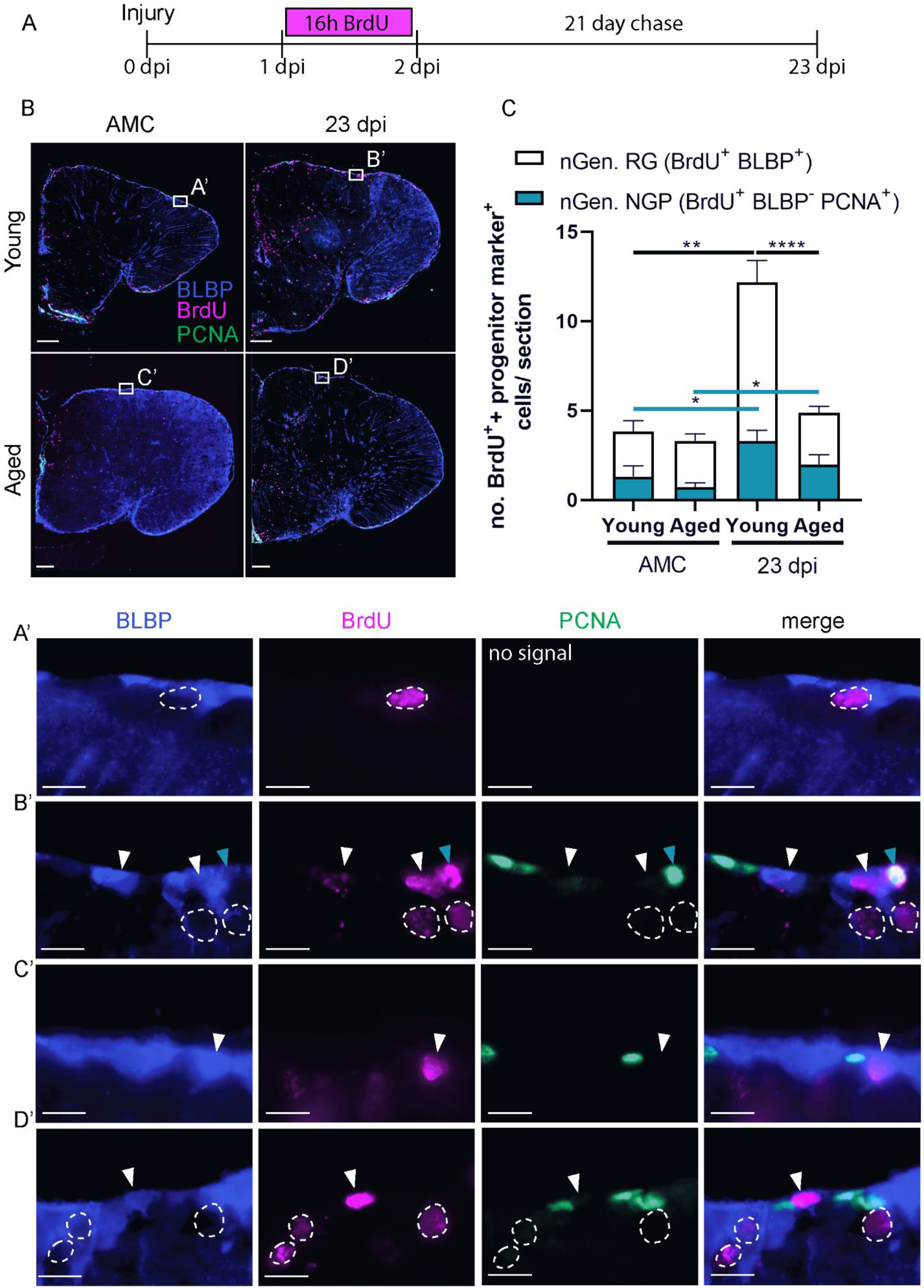
Aging hampers the replenishing of progenitors in the VZ after injury. **(A)** Experimental set up: Day 0: time of a stab-wound injury; From 1 to 2 dpi: injured fish and AMCs are placed in BrdU water for 16 hours: BrdU will be incorporated in the DNA of dividing cells and passed on to the progeny upon each cell division. 23 dpi: regeneration is completed in young fish, all brain samples of young and aged, AMC and injury conditions are collected for cell analysis (as illustrated in B). **(B)** Triple staining for BLBP (blue), BrdU (magenta) and PCNA (green). Boxed areas are magnified in (A’-D’). White arrowheads depict double positive BLBP^+^ BrdU^+^ newly generated RGs. Turquoise arrowheads mark double positive BLBP^-^ BrdU^+^ PCNA^+^ newly generated NGPs. Newly generated RGs, and to a lesser extent newly generated NGPs, are produced after injury in the VZ. Scale bars in (B): 100 µm. Scale bars in (A’-D’): 10 µm. **(C)** Number of BLBP^+^ BrdU^+^ newly generated RGs and BLBP^-^ BrdU^+^ PCNA^+^ newly generated NGPs in the VZ of young and aged telencephali in naive conditions (AMC) and at 23 dpi. Significantly more newly generated progenitors are produced after injury. Young adult fish create significantly more newly generated RGs compared to aged killifish at 23 dpi. This difference is not significant for newly generated NGPs. *p≤0,05, **p≤0,01, ****p≤0,0001; Unpaired t-test or non-parametric Mann Whitney test is used to compare AMC to 23 dpi fish. Two-way ANOVA is used to compare young and aged fish, followed by Sidak’s multiple comparisons test. Values are mean ± SEM; n≥5. RG: radial glia, NGP: non-glial progenitor, AMC: age-matched control, VZ: ventricular zone, dpi: days post injury, nGen: newly generated.

Independent of the progenitor class, we observed that naive fish showed negligible amounts of newborn RG and NGPs. In injured brains, clearly more newly generated progenitors are present in the dorsal VZ, albeit much less in aged fish compared to young adult fish (for newly generated RG: 2,9±0,3636 versus 8,883±1,212; P<0,0001 and for newly generated NGP: 1,967±0,5686 versus 3,3±0,5925; P=n.s.) (Figure 6B,C).

Remarkable, the dividing progenitor pool was replenished to baseline levels during the 21 day chase for each specific age (Figure S5). For both progenitor classes, a similar number of dividing cells was observed compared to their respective AMCs (Figure S5). Aged fish however still had a lower number of dividing NGPs at 23 dpi compared to young adult animals. The number of dividing RGs was very low for both ages.

Taken together, these results indicate that aged killifish are less efficient in replenishing the dorsal VZ after stab-wound injury compared to young adult killifish. Furthermore, generation of neurons and progenitors does not exhaust the available dividing progenitor pools during a short 21 day chase period, suggesting a great capacity of killifish for progenitor cell self-renewal and neuro(re)genesis. This capacity was again higher in young adult killifish.

## Discussion

We designed and benchmarked an easy-to-use and adequate TBI model to study neuroregeneration and uncover the impact of aging on brain repair in the fastest-aging teleost laboratory species, *N. furzeri*. We discovered an age-related decline in proliferation as well as injury-related proliferative response of neurogenic progenitors. The NGP population is most active and responds most prominently to injury. Glial scarring and a strong, localized inflammatory reaction also typify the aged condition post-injury, while young adult brains regenerate in a seamless manner. Taken together, our results provide a validated animal model for future studies to unveil underlying mechanisms driving the loss of full neuroregenerative capacity upon aging.

Stab-wound injuries are reliable, effective injury models and require no specialized equipment, which makes them applicable in a broad range of laboratories. They are also very practical to study a vast array of pathological conditions, since they elicit a clear multicellular read-out, e.g. disruption of the blood brain barrier (BBB), inflammation, astrogliosis. Stab-injuries are often used in mice as TBI model, e.g. Balasingam *et al*., 1994; Allahyari *et al*., 2015; Frik *et al*., 2018; Hashimoto *et al*., 2018; Mattugini *et al*., 2018. They have also been extensively characterized in the zebrafish telencephalon (Ayari *et al*., 2010; Kroehne *et al*., 2011; März *et al*., 2011; Baumgart *et al*., 2012; Kishimoto *et al*., 2012; Kyritsis *et al*., 2012; Barbosa *et al*., 2015). The benefit of translating this model to the killifish is the implementation of the factor age to investigate its influence on the high neuroregenerative capacity, typically associated with teleost species. Surgery can be performed under 5 minutes, no specialized equipment is required and since teleost fish are regenerative-competent vertebrates, full recovery is established within 23-30 days in young adult killifish (data presented here) and adult zebrafish (Ayari *et al*., 2010; Kroehne *et al*., 2011; Kishimoto *et al*., 2012).

The current study provides evidence that the killifish telencephalon is subjected to age-associated stem cell exhaustion, one of the 9 hallmarks of aging (López-Otín *et al*., 2013). Farr less proliferating progenitors, newborn neurons and newly generated progenitors were counted in aged killifish, even after injury, which is in line with other reports in killifish (Tozzini *et al*., 2012) and zebrafish (Edelmann *et al*., 2013; Bhattarai *et al*., 2017). The most striking difference to other teleosts is that the reactive proliferation is related to the atypical NGPs in killifish, instead of the typical RGs that support zebrafish neuroregeneration (Kroehne *et al*., 2011; Baumgart *et al*., 2012). RG division is very low in killifish, even after injury. It was recently discovered that RGs enter a Notch3-dependent quiescence already in larval stages (Coolen *et al*., 2020). Whenever these RGs are reactivated, their division is mainly of gliogenic nature (Coolen *et al*., 2020). The low division of RGs in the killifish brain is reminiscent of adult neural stem cells (NSCs) in the SGZ and SVZ of mice. Here, GFAP^+^ NSCs are mostly quiescent and use predominately asymmetric divisions to create a new NSC and a transient amplifying progenitor. They can however also use symmetric divisions for self-renewal (reviewed in Daynac *et al*., 2017). Indeed, we and others discovered that killifish RGs, are mostly quiescent, have low division potential and can also self-renew (Coolen *et al*., 2020).Whether killifish RGs can also generate NGPs via asymmetric division the way mammalian NSCs create transit amplifying cells remains however elusive. There are some similarities between NGPs and mammalian transient amplifying progenitors. Both progenitor types have a higher division rate compared to NSCs/RGs and are responsible for the production of newborn neurons (neuroblasts) (Daynac *et al*., 2017; Coolen *et al*., 2020). In the SVZ, generation of neurons happen via symmetric divisions, directly creating two neuroblasts from one transit amplifying progenitor (Daynac *et al*., 2017). NGPs on the contrary, also appear to have a lot of self-renewing capacity (Coolen *et al*., 2020). As such, NGPs might represent a separate progenitor lineage, created early in brain development, that is self-sustainable while the RGs become quiescent upon adulthood. In the present study we also discovered that the percentage of dividing RGs among all dividing cells was doubled in aged killifish, although still low in respect to the percentage of dividing NGPs. We hypothesise that when NGPs become exhausted with increasing age, more RGs become activated as a replenishing strategy. This would imply that RGs give rise to NGPs, which remains unexplored to date. Taken together, our results urge for dedicated lineage tracing from these two progenitor pools. Insights into the self-renewal capacities of both progenitor types might unravel key concepts of stem cell biology and division mode necessary for successful neuroregeneration.

Notwithstanding the low percentage of dividing progenitors, aged killifish are still able to produce a neurogenic response upon injury. They generate a considerable number of newborn neurons, albeit much less than young adult killifish. Few neurons reach the injury site in the parenchyma, suggesting migration is also impacted by aging. It remains to be investigated if these neurons are still capable of maturing and integrating in the existing aged circuit, and if some degree of improved functional outcome can be established. The massive surge in microglia/macrophages at the injury site in aged killifish, likely creates chronic inflammation and renders the parenchyma unsuitable for neuron migration, maturation and integration, or may even drive the newborn neurons into apoptosis. This is similar to mammals, in which a large portion of the injury-induced neurons will die due to the pathological environment (Turnley *et al*., 2014; Grade *et al*., 2017). We predict the chronic inflammation to be partly caused by influx of blood-derived macrophages. Aging renders the BBB weak, enabling more macrophages to pass the BBB in aged brains (Montagne *et al*., 2015; Verheggen *et al*., 2020). This will increase the inflammatory response and exacerbate damage. Also in the aged zebrafish telencephalon, increased numbers of ramified, but not round, microglia were discovered after amyloidosis (Bhattarai *et al*., 2017).

Aged killifish also showed glial scarring at the injury site, which is – to our knowledge – never whitnessed before in other regeneration studies working with telencephalic injuries in teleosts (Kroehne *et al*., 2011; Baumgart *et al*., 2012; Edelmann *et al*., 2013; Bhattarai *et al*., 2017). In mammals, the glial scar acts as a physical barrier consisting of reactive astrocytes, NG2 glia and microglia. On the one hand, the glial scar preserves tissue integrity by repairing the BBB and blocking the influx of fibrotic cells and blood-derived macrophages. On the other hand, the scar represents a physical barrier for axonal outgrowth, thereby restricting neural circuit integration and overall brain repair (reviewed in Adams *et al*., 2018). We hypothesize that also in aged injured killifish, the glial scar is preventing newborn neurons to integrate and thereby impedes successful brain recovery. Interestingly, the presence of RG fibers, expressing high amounts of GS, suggests a function of RGs in glial scar formation. GS is known to convert glutamate into glutamine, reducing toxic extracellular glutamate levels (Zou *et al*., 2010). Its presence at the glial scar could indicate that high glutamate toxicity levels were still present in aged killifish. The cells surrounding the scar tissue indeed had a swollen morphology, typical for cytotoxicity (Liang *et al*., 2007).

Since our data predict that RGs do not support the production of new neurons in the killifish brain, the question remains what role RGs play after injury. At 2 dpi, RG fibers appeared swollen in aged injured killifish, suggesting that the RGs do respond to the insult. Considering that many of the mammalian astrocyte-specific genes (GLT-1, BLBP, GS, GLAST, aldh1l1, GFAP, Vimentin, S100β) are also expressed by teleost RGs (Ganz *et al*., 2010; März *et al*., 2010; Chen *et al*., 2020; Coolen *et al*., 2020), a similar function is highly likely. In mammals, RGs act as neural stem cells early in development and support neurogenesis. Later on, most RGs leave the ventricular zone – the mammalian stem cell zone – and become star-shaped astrocytes (reviewed in Kriegstein *et al*., 2009). ‘Star-shaped astroglia’ have previously been described in the zebrafish spinal cord (Kawai *et al*., 2001), but are are yet to be found in the teleost telencephalon. New evidence was recently provided that teleost ‘astroglia’ resemble mammalian astrocytes more than once thought. In the larval zebrafish spinal cord they are in close association with synapses, exhibit tiling and calcium signaling dynamics and have a long bushy morphology that expands during development (Chen *et al*., 2020). In the zebrafish larval brain, interplay between astroglia and neurons during epileptic seizures was discovered, as well as large gap junction-coupled glial networks (Diaz Verdugo *et al*., 2019). It thus seems that teleost RGs keep their developmental morphology into adulthood, but adopt several functions typically associated with mammalian protoplasmic astrocytes (Wahis *et al*., 2021). This might also explain why teleosts have such impressive regenerative abilities as their astroglia resemble more the ‘developmental’ mammalian RGs, which makes them highly neurogenic into adulthood. The killifish thus represents a promising vertebrate model to unravel the function of the teleost ‘astroglia’ because the delineation between RG stem cell properties and astrocyte-properties seems more distinguished than in other teleost models. How these killifish ‘astroglia’ respond to injury and, interestingly, behave upon brain aging, are intriguing research questions for the future.

Altogether, we expose the aged killifish to model low regenerative abilities similar to adult mammals. The aged neuroregenerative response is characterized by glial scarring, secondary damage, aggravating inflammation, reduced proliferation of stem cells and reduced production of new neurons. Our model will therefore be highly useful to elucidate how to reverse the aging state of the brain in order to reinstate a high neuroreparative strategy as occruing in young adult killifish. Such insights will hopefully play a pivotal role in boosting neurorepair in mammals in the near future.

## Author Contributions

**J.V.H**.: Conceptualization, design, experiments, statistical analysis, writing, original draft, review, editing and visualization. **V.M**.: Experiments, statistical analysis, review and editing. **S.V**.: Experiments. **C.Z**., **L.M**., **R.A**., **E.S**.: Review and editing. **L.A**.: Conceptualization, design, writing, review, editing, study supervision.

## List of abbreviations

AMC: age-matched control
BBB: brain blood barrier
BLBP: brain lipid-binding protein
CNS: central nervous system
dpi: days post injury
GFAP: glial fibrillary acidic protein
GS: glutamine synthetase
hpi: hours post injury
NGP: non-glial progenitor
NPC: neuronal progenitor cell
NSC: neuronal stem cell
PCNA: proliferating cell nuclear antigen
PVZ: periventricular zone
RG: radial glia
RMS: rostral migratory stream
SGZ: subgranular zone
SOX2: SRY-box transcription factor 2
SVZ: subventicular zone
TBI: traumatic brain injury
TUNEL: terminal deoxynucleotidyl transferase-mediated dUTP nick end labeling
VZ: ventricular zone

## Acknowledgments

We thank our collaborators of the KU Leuven killifish consortium Prof. Dr. L. Brendonck and Dr. T. Pinceel to provide African turquoise killifish eggs of the short-lived GRZ-AD strain. We are grateful to Dr. T. Pinceel for his excellent problem-solving skills in relation to killifish breeding and housing. We thank Rony Van Aerschot, Arnold Van Den Eynde, and the KU Leuven killifish team for taking care of the animals and setting up our shared fish facility. We are thankful to Ria Vanlaer and Lieve Geenen for their technical support. Students that were involved in optimizing techniques are Niels Vidal, Karen Libberecht and Lotte De Bauw. We are also grateful to Dr. Jessie Van houcke, Dr. S. Gilissen and Dr. M. Hennes for reviewing this manuscript. We acknowledge Biorender.com for delivering the tools to create Figure S4.

## Supplementary figure legends

**Supplementary figure 1: Research methodology and glial scarring**

**(A)** Survival curve of *N. furzeri* females (strain GRZ-AD), housed in a Tecniplast ZebTEC multi-linking aquarium system (n=48). 6 week-old killifish are chosen as young adult based on 100% survival rate and having reached sexual maturity. 18 week-old killifish are chosen as aged based on 76,6% survival rate and showing phenotypic aging hallmarks (illustrated in B).

**(B)** Photographs of a young adult (6 week-old) and aged (18 week-old) *N. furzeri* females (strain GRZ-AD). Aged females show phenotypic aging hallmarks, such as a spinal curvature and protrusion of the lip (arrowheads).

**(C)** Schematic of the stab-wound injury method. After removing skin above the skull, a 33-gauge Hamiliton needle is pushed into the medial zone of the right telencephalic hemisphere. The needle is dipped in DiD solution (red), which labels the membranes of cells around the injury site (illustrated in D,E).

**(D-K)** The injury site can be found by scanning sections for DiD-positivity (red crystals) after cryosectioning, for example at 30 dpi. (D,F,H, J) Adjacent coronal sections of a young adult killifish telencephalon. (D) The DiD dye is still visible at 30 dpi. (F) Cresyl violet staining shows that the tissue of the young adult fish is structurally regenerated. (H) No glial scar (or malformation) is visible after staining for L-plastin^+^ microglia/macrophages in the young adult brain at the site of injury. (J) Gs^+^ RG fiber distribution is normal in young adult fish at 30 dpi.

**(E**,**G**,**I)** Adjacent coronal sections of an aged killifish telencephalon. (E) The DiD dye clearly marks the injury site at 30 dpi. (G) Cresyl violet staining shows tissue scarring (arrowhead in G’), indicative for incomplete repair. (I) Signs of glial scarring are visible after staining for L-plastin^+^ microglia/macrophages in the aged brain at the site of injury. (K) Gs^+^ RG fibers are in close contact with the glial scar (arrowhead in K’).

(D’,E’,F’,G’,H’,I’) are magnifications of the boxed area in (D,E,F,G,H,I) respectively. Scale bars in D-I: 100 µm. Scale bars in D’-I’: 50 µm. dpi: days post injury.

**Supplementary figure 2: Methodology of injury surface area measurements**

In ImageJ (FIJI) the scale is set and a polygon is drawn around the injury site with the polygon tool (blue line). ImageJ calculates the surface of the polygon. At early stages, the injury site is visible by a blood clot and the borders of this clot are taken as the injury borders (young, aged at 1 dpi, scale bars: 50 µm). In later stages, only a malformation of the parenchymal tissue is visible (aged 30 dpi, scale bar: 50 µm). In this case, the borders of this malformation are taken as the injury borders (Illustrated in the inset of aged 30 dpi, scale bar: 20 µm).

**Supplementary figure 3: Injury induces proliferation of microglia in young and aged fish**

**(A)** Double staining for L-plastin (magenta) and PCNA (green) with DAPI (blue) shows proliferating microglia/macrophages (arrowhead) on coronal sections of young and aged killifish at 2 dpi. Scale bars: 10 µm.

**(B)** Absolute number of double positive L-plastin^+^ PCNA^+^ proliferating microglia/macrophages in young adult and aged telencephali in naive conditions and at 1 and 2 dpi. Microglia/macrophages start proliferating early after injury in the killifish telencephalon. Significantly higher levels of proliferating microglia/macrophages are observed at 1 and 2 dpi for both ages. This activation is slightly more pronounced but highly variable in aged killifish compared to young adult fish. *p≤0,05, **p≤0,01; One-way ANOVA is used to compare naive fish to injured fish for each. Young: parametric one-way ANOVA, followed by Dunnett’s multiple comparisons test. Aged: non-parametric Kruskal-Wallis test, followed by Dunn’s multiple comparisons test. Two-way ANOVA is used to compare young and aged fish, followed by Sidak’s multiple comparisons test. Values are mean ± SEM; n≥5.

**Supplementary figure 4: Counting method**

Schematic view of the different regions in which different types of cells are counted in the right telencephalic hemisphere. VZ: ventricular zone, PVZ: periventricular zone, RMS: rostral migratory stream, RG: radial glia, NGP: non-glial progenitor. Created with BioRender.com.

**Supplementary table 1: Percentages of dividing RGs and NGPs among the total amount of dividing progenitors differ between young adult and aged killifish**

Independent of injury, aged killifish display more dividing RGs in regard to the total amount of dividing progenitors compared to young adult fish. *p≤0,05; Unpaired T-test is used to compare the percentage of dividing RGs or NGPs among all dividing progenitors between young adult and aged killifish. Values are mean ± SEM; n≥5.

**Supplementary figure 5: The dividing progenitor pool is not depleted within a short period after injury**

**(A)** Absolute number of BLBP^+^ PCNA^+^ dividing RGs and **(B)** BLBP^-^ PCNA^+^ dividing NGPs in the VZ of young and aged telencephali in naive conditions (AMC) and at 23 dpi. Even after the production of neurons, the number of dividing RGs and NGPs is respectively similar or increased at 23 dpi compared to AMCs. This suggests that killifish replenish the proliferative stem cell pool after injury. *p≤0,05, **p≤0,01, Unpaired t-test or non-parametric Mann Whitney test is used to compare AMC to 23 dpi fish. Two-way ANOVA is used to compare young and aged fish, followed by Sidak’s multiple comparisons test. Values are mean ± SEM; n≥5. RG: radial glia, NGP: non-glial progenitor, AMC: age-matched control, VZ: ventricular zone, dpi: days post injury.

## References

Adams, K. L. and Gallo, V. (2018) The diversity and disparity of the glial scar, Nature Neuroscience, 21(1), pp. 9–15. doi: 10.1038/s41593-017-0033-9.

Adolf, B., Chapouton, P., Lam, C. S., Topp, S., Tannhäuser, B., Strähle, U., et al. (2006) Conserved and acquired features of adult neurogenesis in the zebrafish telencephalon, Developmental Biology, 295(1), pp. 278–293. doi: 10.1016/j.ydbio.2006.03.023.

Allahyari, R. V. and Garcia, A. D. R. (2015) Triggering Reactive Gliosis In Vivo by a Forebrain Stab Injury, Journal of Visualized Experiments, (100), p. e52825. doi: 10.3791/52825.

Arvidsson, A., Collin, T., Kirik, D., Kokaia, Z. and Lindvall, O. (2002) Neuronal replacement from endogenous precursors in the adult brain after stroke, Nature Medicine, 8(9), pp. 963–70. doi: 10.1038/nm747.

Ayari, B., El Hachimi, K. H., Yanicostas, C., Landoulsi, A. and Soussi-Yanicostas, N. (2010) Prokineticin 2 expression is associated with neural repair of injured adult zebrafish telencephalon, Journal of Neurotrauma, 27(5), pp. 959–72. doi: 10.1089/neu.2009.0972.

Balasingam, V., Tejada-Berges, T., Wright, E., Bouckova, R. and Yong, V. (1994) Reactive astrogliosis in the neonatal mouse brain and its modulation by cytokines, The Journal of Neuroscience, 14(2), pp. 846–856. doi: 10.1523/JNEUROSCI.14-02-00846.1994.

Barbosa, J. S., Sanchez-Gonzalez, R., Di Giaimo, R., Baumgart, E. V., Theis, F. J., Götz, M., et al. (2015) Live imaging of adult neural stem cell behavior in the intact and injured zebrafish brain, Science, 348(6236), pp. 789–793. doi: 10.1126/science.aaa2729.

Baumgart, E. V, Barbosa, J. S., Bally-Cuif, L., Gotz, M. and Ninkovic, J. (2012) Stab wound injury of the zebrafish telencephalon: A model for comparative analysis of reactive gliosis, Glia, 60(3), pp. 343–357. doi: 10.1002/glia.22269.

Bhattarai, P., Thomas, A. K., Zhang, Y. and Kizil, C. (2017) The effects of aging on Amyloid-beta42-induced neurodegeneration and regeneration in adult zebrafish brain, Neurogenesis (Austin), 4(1), p. e1322666. doi: 10.1080/23262133.2017.1322666.

Chen, J., Poskanzer, K. E., Freeman, M. R. and Monk, K. R. (2020) Live-imaging of astrocyte morphogenesis and function in zebrafish neural circuits, Nature Neuroscience, 23, pp. 1297– 1306. doi: 10.1038/s41593-020-0703-x.

Coolen, M., Labusch, M., Mannioui, A. and Bally-Cuif, L. (2020) Mosaic Heterochrony in Neural Progenitors Sustains Accelerated Brain Growth and Neurogenesis in the Juvenile Killifish N. furzeri, Current Biology, 30(4), pp. 736-745.e4. doi: 10.1016/j.cub.2019.12.046.

Daynac, M. and Petritsch, C. K. (2017) Regulation of Asymmetric Cell Division in Mammalian Neural Stem and Cancer Precursor Cells, in Tassan, J. P.and Kubiak, J. (eds) Asymmetric Cell Division in Development, Differentiation and Cancer. Results and Problems in Cell Differentiation. vol. 61. Springer, Cham, pp. 375–399. doi: 10.1007/978-3-319-53150-2_17.

Diaz Verdugo, C., Myren-Svelstad, S., Aydin, E., Van Hoeymissen, E., Deneubourg, C., Vanderhaeghe, S., et al. (2019) Glia-neuron interactions underlie state transitions to generalized seizures, Nature Communications, 10(3830), pp. 1–13. doi: 10.1038/s41467-019-11739-z.

Ding, L., Kuhne, W. W., Hinton, D. E., Song, J. and Dynan, W. S. (2010) Quantifiable biomarkers of normal aging in the Japanese Medaka fish (Oryzias latipes), PLoS ONE, 5(10), p. e13287. doi: 10.1371/journal.pone.0013287.

Dugger, B. N. and Dickson, D. W. (2017) Pathology of Neurodegenerative Diseases, Cold Spring Harbor Perspectives in Biology, 9(7), p. a028035. doi: 10.1101/cshperspect.a028035.

Edelmann, K., Glashauser, L., Sprungala, S., Hesl, B., Fritschle, M., Ninkovic, J., et al. (2013) Increased radial glia quiescence, decreased reactivation upon injury and unaltered neuroblast behavior underlie decreased neurogenesis in the aging zebrafish telencephalon, J Comp Neurol, 521(13), pp. 3099–3115. doi: 10.1002/cne.23347.

El-Hayek, Y. H., Wiley, R. E., Khoury, C. P., Daya, R. P., Ballard, C., Evans, A. R., et al. (2019) Tip of the Iceberg: Assessing the Global Socioeconomic Costs of Alzheimer’s Disease and Related Dementias and Strategic Implications for Stakeholders, Journal of Alzheimer’s Disease, 70(2), pp. 323–341. doi: 10.3233/JAD-190426.

Frik, J., Merl-Pham, J., Plesnila, N., Mattugini, N., Kjell, J., Kraska, J., et al. (2018) Cross-talk between monocyte invasion and astrocyte proliferation regulates scarring in brain injury, EMBO reports, 19(5). doi: 10.15252/embr.201745294.

Galvan, V. and Jin, K. (2007) Neurogenesis in the aging brain., Clinical interventions in aging, 2(4), pp. 605–610. doi: 10.2147/cia.s1614.

Ganz, J., Kaslin, J., Hochmann, S., Freudenreich, D. and Brand, M. (2010) Heterogeneity and Fgf dependence of adult neural progenitors in the zebrafish telencephalon, GLIA, 58(11), pp. 1345– 63. doi: 10.1002/glia.21012.

Genade, T., Benedetti, M., Terzibasi, E., Roncaglia, P., Valenzano, D. R., Cattaneo, A., et al. (2005) Annual fishes of the genus Nothobranchius as a model system for aging research, Aging Cell, 4(5), pp. 223–233. doi: 10.1111/j.1474-9726.2005.00165.x.

Gerhard, G. S., Kauffman, E. J., Wang, X., Stewart, R., Moore, J. L., Kasales, C. J., et al. (2002) Life spans and senescent phenotypes in two strains of Zebrafish (Danio rerio), Experimental Gerontology, 37(8–9), pp. 1055–68. doi: 10.1016/S0531-5565(02)00088-8.

Gopalakrishnan, S., Cheung, N. K. M., Yip, B. W. P. and Au, D. W. T. (2013) Medaka fish exhibits longevity gender gap, a natural drop in estrogen and telomere shortening during aging: A unique model for studying sex-dependent longevity, Frontiers in Zoology, 10(1), p. 78. doi: 10.1186/1742-9994-10-78.

Grade, S. and Götz, M. (2017) Neuronal replacement therapy: previous achievements and challenges ahead, npj Regenerative Medicine, 2, p. 29. doi: 10.1038/s41536-017-0033-0.

Grandel, H., Kaslin, J., Ganz, J., Wenzel, I. and Brand, M. (2006) Neural stem cells and neurogenesis in the adult zebrafish brain: Origin, proliferation dynamics, migration and cell fate, Developmental Biology, 295(1), pp. 263–277. doi: 10.1016/j.ydbio.2006.03.040.

Grupp, L., Wolburg, H. and Mack, A. F. (2010) Astroglial structures in the zebrafish brain, Journal of Comparative Neurology. doi: 10.1002/cne.22481.

Hashimoto, K., Nakashima, M., Hamano, A., Gotoh, M., Ikeshima-Kataoka, H., Murakami-Murofushi, K., et al. (2018) 2-carba cyclic phosphatidic acid suppresses inflammation via regulation of microglial polarisation in the stab-wounded mouse cerebral cortex, Scientific Reports, 8(1), p. 9715. doi: 10.1038/s41598-018-27990-1.

Hou, Y., Dan, X., Babbar, M., Wei, Y., Hasselbalch, S. G., Croteau, D. L., et al. (2019) Ageing as a risk factor for neurodegenerative disease, Nature Reviews Neurology, 15(10), pp. 565–581. doi: 10.1038/s41582-019-0244-7.

Howe, K., Clark, M. D., Torroja, C. F., Torrance, J., Berthelot, C., Muffato, M., et al. (2013) The zebrafish reference genome sequence and its relationship to the human genome, Nature. 2013/04/19, 496(7446), pp. 498–503. doi: 10.1038/nature12111.

Hu, C.-K. and Brunet, A. (2018) The African turquoise killifish: A research organism to study vertebrate aging and diapause, Aging Cell, 17(3), p. e12757. doi: 10.1111/acel.12757.

Kawai, H., Arata, N. and Nakayasu, H. (2001) Three-dimensional distribution of astrocytes in zebrafish spinal cord, GLIA, 36, pp. 406–413. doi: 10.1002/glia.1126.

Kernie, S. G. and Parent, J. M. (2010) Forebrain neurogenesis after focal Ischemic and traumatic brain injury, Neurobiol Dis. 2009/11/17, 37(2), pp. 267–274. doi: 10.1016/j.nbd.2009.11.002.

Kim, C. H., Ueshima, E., Muraoka, O., Tanaka, H., Yeo, S. Y., Huh, T. L., et al. (1996) Zebrafish elav/HuC homologue as a very early neuronal marker, Neuroscience Letters, 216(2), pp. 109–112. doi: 10.1016/0304-3940(96)13021-4.

Kim, Y., Nam, H. G. and Valenzano, D. R. (2016) The short-lived African turquoise killifish: An emerging experimental model for ageing, Disease Models and Mechanisms, 9(2), pp. 115–129.

Kishimoto, N., Shimizu, K. and Sawamoto, K. (2012) Neuronal regeneration in a zebrafish model of adult brain injury, Dis Model Mech. 2011/10/27, 5(2), pp. 200–209. doi: 10.1242/dmm.007336.

Kriegstein, A. and Alvarez-Buylla, A. (2009) The Glial Nature of Embryonic and Adult Neural Stem Cells, Annual Review of Neuroscience, 32(1), pp. 149–184. doi: 10.1146/annurev.neuro.051508.135600.

Kroehne, V., Freudenreich, D., Hans, S., Kaslin, J. and Brand, M. (2011) Regeneration of the adult zebrafish brain from neurogenic radial glia-type progenitors, Development, 138(22), pp. 4831– 4841. doi: 10.1242/dev.072587.

Kyritsis, N., Kizil, C., Zocher, S., Kroehne, V., Kaslin, J., Freudenreich, D., et al. (2012) Acute inflammation initiates the regenerative response in the adult zebrafish brain, Science, 338(6112), pp. 1353–6. doi: 10.1126/science.1228773.

Liang, D., Bhatta, S., Gerzanich, V. and Simard, J. M. (2007) Cytotoxic edema: mechanisms of pathological cell swelling, Neurosurgical Focus, 22(5), pp. 1–9. doi: 10.3171/foc.2007.22.5.3.

López-Otín, C., Blasco, M. A., Partridge, L., Serrano, M. and Kroemer, G. (2013) The Hallmarks of Aging, Cell, 153(6), pp. 1194–1217. doi: 10.1016/j.cell.2013.05.039.

Marques, I. J., Lupi, E. and Mercader, N. (2019) Model systems for regeneration: Zebrafish, Development (Cambridge), 146(18), pp. 1–13. doi: 10.1242/dev.167692.

März, M., Chapouton, P., Diotel, N., Vaillant, C., Hesl, B., Takamiya, M., et al. (2010) Heterogeneity in progenitor cell subtypes in the ventricular zone of the zebrafish adult telencephalon, GLIA, 58(7), pp. 870–88. doi: 10.1002/glia.20971.

März, M., Schmidt, R., Rastegar, S. and Strahle, U. (2011) Regenerative response following stab injury in the adult zebrafish telencephalon, Developmental Dynamics, 240(9), pp. 2221–31. doi: 10.1002/dvdy.22710.

Mattugini, N., Merl-Pham, J., Petrozziello, E., Schindler, L., Bernhagen, J., Hauck, S. M., et al. (2018) Influence of white matter injury on gray matter reactive gliosis upon stab wound in the adult murine cerebral cortex, Glia, 66(8), pp. 1644–1662. doi: 10.1002/glia.23329.

Miller, R. A., Harper, J. M., Dysko, R. C., Durkee, S. J. and Austad, S. N. (2002) Longer life spans and delayed maturation in wild-derived mice, Experimental Biology and Medicine, 227(7), pp. 500–8. doi: 10.1177/153537020222700715.

Montagne, A., Barnes, S. R., Sweeney, M. D., Halliday, M. R., Sagare, A. P., Zhao, Z., et al. (2015) Blood-Brain barrier breakdown in the aging human hippocampus, Neuron. doi: 10.1016/j.neuron.2014.12.032.

Mueller, T., Dong, Z., Berberoglu, M. A. and Guo, S. (2011) The dorsal pallium in zebrafish, Danio rerio (Cyprinidae, Teleostei), Brain Research. doi: 10.1016/j.brainres.2010.12.089.

Mueller, T. and Wullimann, M. F. (2009) An evolutionary interpretation of teleostean forebrain anatomy, in Brain, Behavior and Evolution. doi: 10.1159/000229011.

Platzer, M. and Englert, C. (2016) Nothobranchius furzeri: A Model for Aging Research and More, Trends Genet. 2016/07/19, 32(9), pp. 543–552. doi: 10.1016/j.tig.2016.06.006.

Polačik, M., Blažek, R. and Reichard, M. (2016) Laboratory breeding of the short-lived annual killifish Nothobranchius furzeri, Nature Protocols, 11, pp. 1396–1413. doi: 10.1038/nprot.2016.080.

Popa-Wagner, A., Buga, A. M. and Kokaia, Z. (2011) Perturbed cellular response to brain injury during aging, Ageing Research Reviews. doi: 10.1016/j.arr.2009.10.008.

Reichwald, K., Petzold, A., Koch, P., Downie, B. r, Hartmann, N., Pietsch, S., et al. (2015) Insights into Sex Chromosome Evolution and Aging from the Genome of a Short-Lived Fish, Cell, 163(6), pp. 1527–1538. doi: 10.1016/j.cell.2015.10.071.

Rothenaigner, I., Krecsmarik, M., Hayes, J. A., Bahn, B., Lepier, A., Fortin, G., et al. (2011) Clonal analysis by distinct viral vectors identifies bona fide neural stem cells in the adult zebrafish telencephalon and characterizes their division properties and fate, Development, 138(8), pp. 1459–1469. doi: 10.1242/dev.058156.

Tanaka, E. M. and Ferretti, P. (2009) Considering the evolution of regeneration in the central nervous system, Nature Reviews Neuroscience, 10(10), pp. 713–723. doi: 10.1038/nrn2707.

Thored, P., Arvidsson, A., Cacci, E., Ahlenius, H., Kallur, T., Darsalia, V., et al. (2006) Persistent Production of Neurons from Adult Brain Stem Cells During Recovery after Stroke, Stem Cells, 24(3), pp. 739–747. doi: 10.1634/stemcells.2005-0281.

Tozzini, E. T., Baumgart, M., Battistoni, G. and Cellerino, A. (2012) Adult neurogenesis in the short-lived teleost Nothobranchius furzeri: localization of neurogenic niches, molecular characterization and effects of aging, Aging Cell, 11(2), pp. 241–251. doi: 10.1111/j.1474-9726.2011.00781.x.

Turnley, A. M., Basrai, H. S. and Christie, K. J. (2014) Is integration and survival of newborn neurons the bottleneck for effective neural repair by endogenous neural precursor cells?, Frontiers in Neuroscience, 8. doi: 10.3389/fnins.2014.00029.

Valdesalici, S. and Cellerino, A. (2003) Extremely short lifespan in the annual fish Nothobranchius furzeri, Proceedings of the Royal Society of London. Series B: Biological Sciences, 270(suppl_2), pp. S189–S191. doi: 10.1098/rsbl.2003.0048.

Valenzano, D. R., Benayoun, B. A., Singh, P. P., Zhang, E., Etter, P. D., Hu, C.-K., et al. (2015) The African Turquoise Killifish Genome Provides Insights into Evolution and Genetic Architecture of Lifespan., Cell, 163(6), pp. 1539–1554. doi: 10.1016/j.cell.2015.11.008.

Van houcke, J., De Groef, L., Dekeyster, E. and Moons, L. (2015) The zebrafish as a gerontology model in nervous system aging, disease, and repair, Ageing Research Reviews, 24(Pt B), pp. 358– 368. doi: 10.1016/j.arr.2015.10.004.

Van houcke, J., Mariën, V., Zandecki, C., Seuntjens, E., Ayana, R. and Arckens, L. (2021) Modeling neuroregeneration and neurorepair in an aging context: the power of a teleost model, Front. Cell Dev. Biol., accepted. doi: 10.3389/fcell.2021.619197.

Verheggen, I. C. M., de Jong, J. J. A., van Boxtel, M. P. J., Gronenschild, E. H. B. M., Palm, W. M., Postma, A. A., et al. (2020) Increase in blood–brain barrier leakage in healthy, older adults, GeroScience. doi: 10.1007/s11357-020-00211-2.

Wahis, J., Hennes, M., Arckens, L. and Holt, M. G. (2021) Star power: the emerging role of astrocytes as neuronal partners during cortical plasticity, Current Opinion in Neurobiology, 67, pp. 174–182. doi: 10.1016/j.conb.2020.12.001.

Wendler, S., Hartmann, N., Hoppe, B. and Englert, C. (2015) Age-dependent decline in fin regenerative capacity in the short-lived fish Nothobranchius furzeri, Aging Cell, 14(5), pp. 857– 866. doi: 10.1111/acel.12367.

Zambusi, A. and Ninkovic, J. (2020) Regeneration of the central nervous system-principles from brain regeneration in adult zebrafish, World Journal of Stem Cells, 12(1), pp. 8–24. doi: 10.4252/wjsc.v12.i1.8.

Zhao, A., Qin, H. and Fu, X. (2016) What determines the regenerative capacity in animals?, BioScience, 66(9), pp. 735–746. doi: 10.1093/biosci/biw079.

Zou, J., Wang, Y.-X., Dou, F.-F., Lü, H.-Z., Ma, Z.-W., Lu, P.-H., et al. (2010) Glutamine synthetase down-regulation reduces astrocyte protection against glutamate excitotoxicity to neurons, Neurochemistry International, 56(4), pp. 577–584. doi: 10.1016/j.neuint.2009.12.021.

Zupanc, G. K. H. (2001) Adult Neurogenesis and Neuronal Regeneration in the Central Nervous System of Teleost Fish, Brain, Behavior and Evolution, 58(5), pp. 250–275. doi: 10.1159/000057569.

Zupanc, G. K. H. and Sîrbulescu, R. F. (2012) Teleost Fish as a Model System to Study Successful Regeneration of the Central Nervous System, in Heber-Katz, E.and Stocum, D. (eds) New Perspectives in Regeneration. Current Topics in Microbiology and Immunology. vol 367. Springer, Berlin, Heidelberg, pp. 193–233. doi: 10.1007/82_2012_297.

