## Supplementary figures for "Aging impairs the essential contributions of non-glial progenitors to neurorepair in the dorsal telencephalon of the Killifish *N. furzeri*"

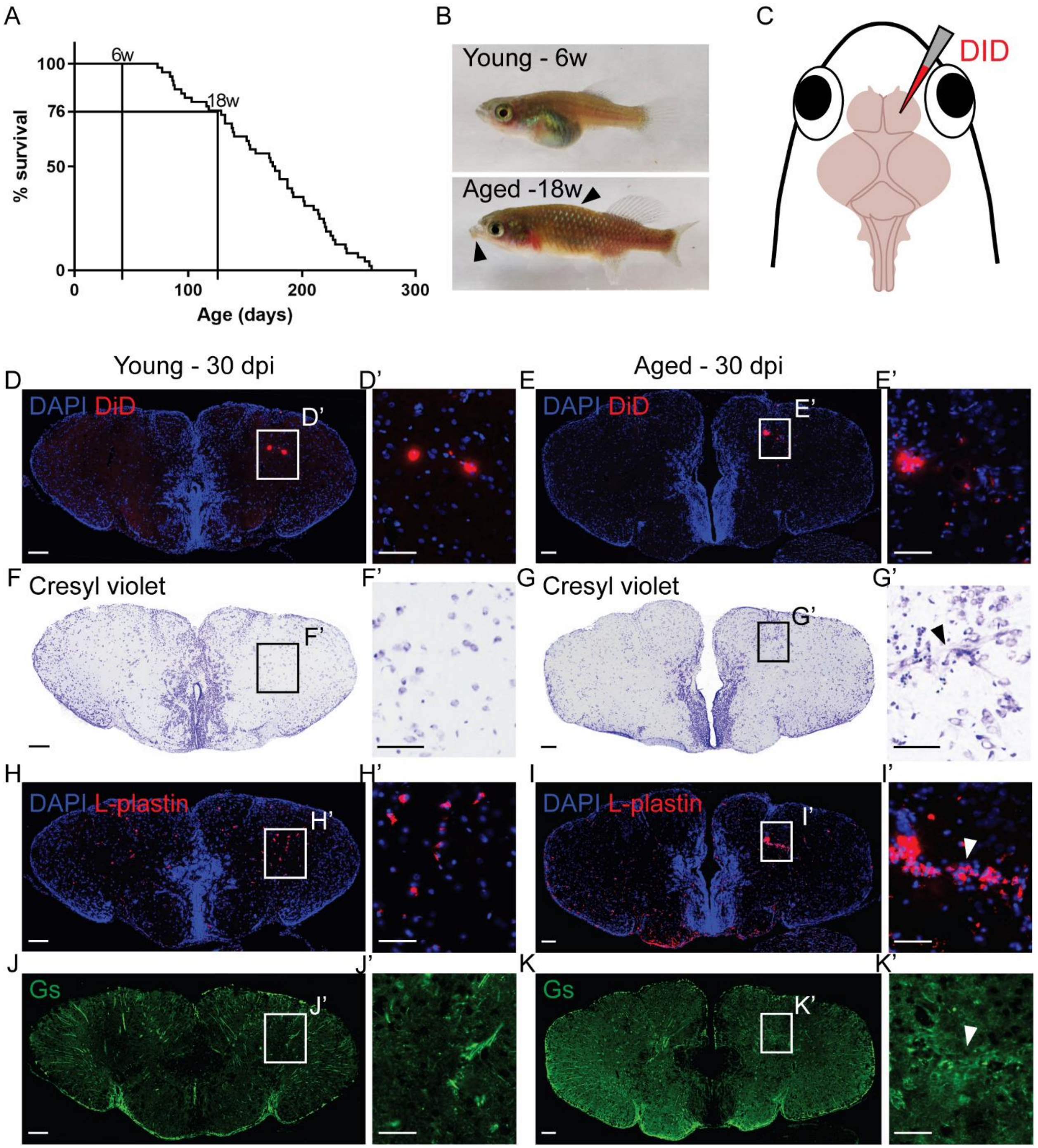


Supplementary figure 1

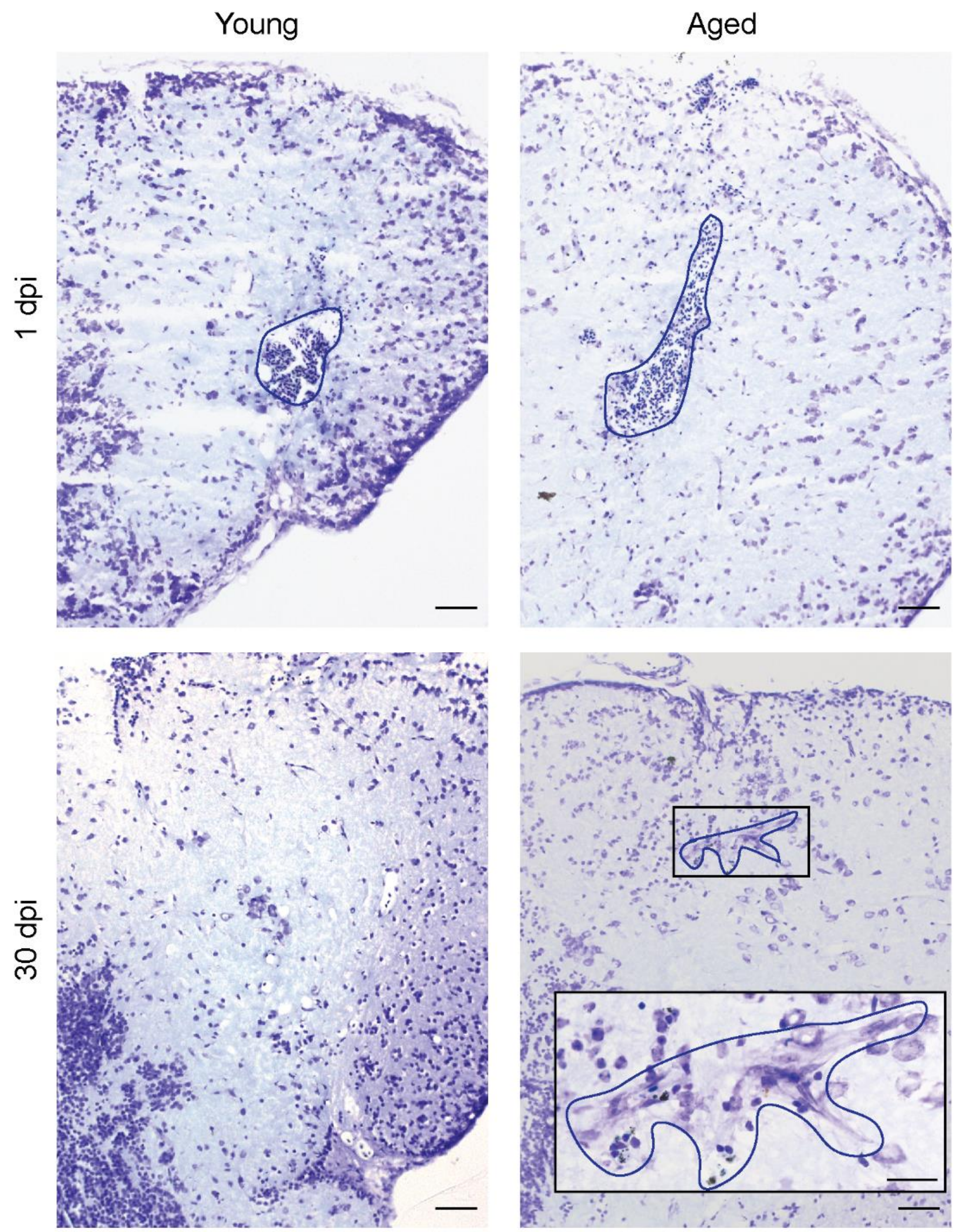


Supplementary figure 2

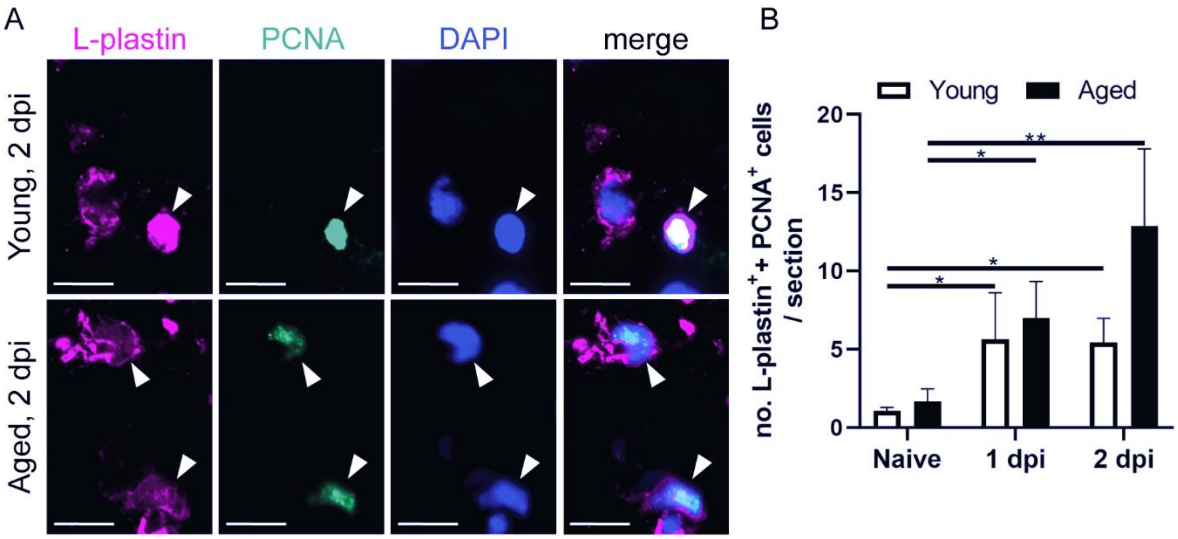


Supplementary figure 3

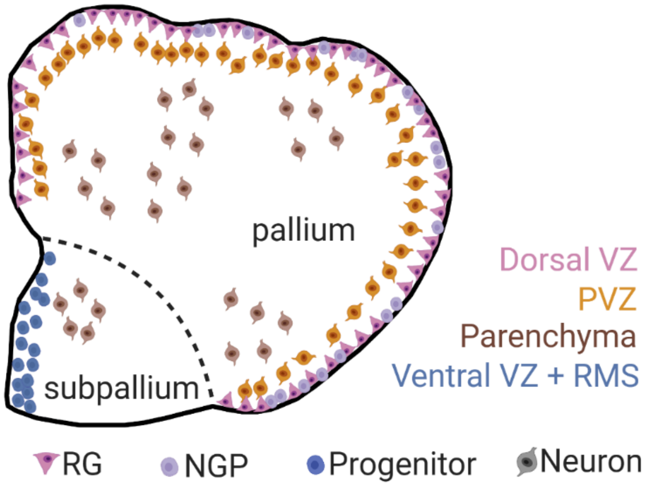


Supplementary figure 4

Supplementary table 1

| | Naive | | | 2 dpi | | |
|---|---|---|---|---|---|---|
| | Young adult | Aged | P-value | Young adult | Aged | P-value |
| **% Dividing RGs** | 6,683% ±1,158 | 17,24% ±3,286 | *P= 0,0163 | 6,509% ±0,688 | 12,28% ±1,837 | *P= 0,0237 |
| **% Dividing NGPs** | 93,32% ±1,158 | 82,76% ±3,286 | | 93,49% ±0,688 | 87,72% ±1,837 | |

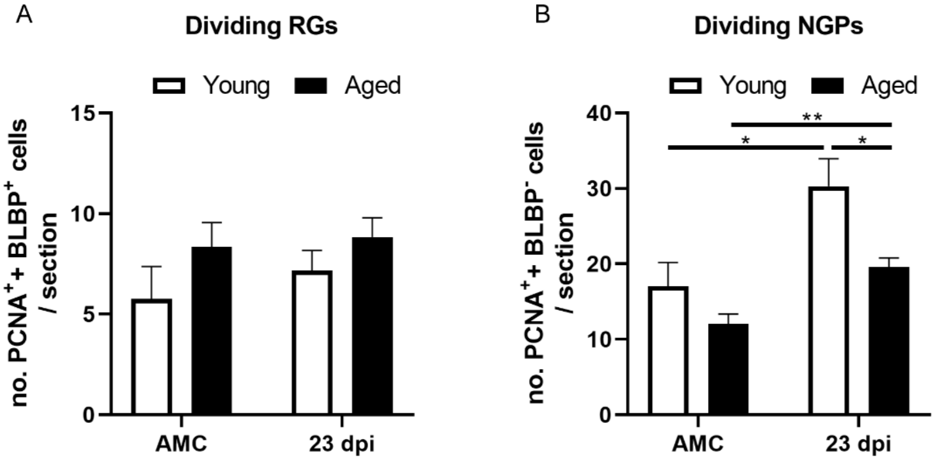


Supplementary figure 5
